# Single-cell analysis reveals cellular heterogeneity and limits of marker-based assessment in retinal ganglion cell-enriched organoid cultures

**DOI:** 10.64898/2026.02.08.704697

**Authors:** Jessica Yuen Wuen Ma, Dulce B. Vargas-Landin, Janya Grainok, Alice Pébay

## Abstract

Human pluripotent stem cell (hPSC)-derived retinal organoids provide an *in vitro* system for generating retinal ganglion cells (RGCs), yet the cellular composition and developmental fidelity of RGC-enriched cultures remain insufficiently characterised. Here, we tested an RGC-enriched approach involving dissociation of hPSC-derived retinal organoids at day 40, corresponding to peak expression of RGC markers, followed by two-dimensional culture conditions intended to enrich for RGC survival. Flow cytometry was used to assess the expression of RGC markers, including POU4F, ISL1, SNCG, and THY1. Across four samples, POU4F expression ranged from 79-95%, ISL1 from 18-58%, SNCG from 22%-91% and THY1 from 3%-29%, indicating substantial variability between markers and samples. Single-cell RNA sequencing analysis of 73,642 cells identified multiple retinal lineages, including retinal progenitors, RGCs, photoreceptor-committed cells, amacrine and horizontal cells, and retinal pigment epithelium (RPE), as well as off-target populations comprising *HOX*-enriched posterior neural cells and other cell types. Cellular composition varied across samples. Transcriptomically defined RGCs accounted for 19-45% of cells across samples, with different subtypes identified. These findings indicate that marker-based assessments alone may overestimate RGC identity and provide a detailed single-cell characterisation of cellular heterogeneity in RGC-enriched retinal organoid cultures.

## Introduction

The human retina offers a unique and accessible window into the organisation of the central nervous system. Retinal ganglion cells (RGCs), the sole projection neurons of the retina, convey visual information to central targets via the optic nerve. Loss of RGCs is a defining pathological feature of optic neuropathies such as glaucoma, which remains a leading cause of irreversible blindness worldwide [1]. Current therapies slow disease progression by lowering intraocular pressure but cannot restore RGCs once degenerated [2], highlighting a critical need for physiologically relevant human models capable of recapitulating RGC development, maturation, and vulnerability to support disease modelling, drug screening, and potential cell-based replacement therapies. However, achieving this goal requires reliable access to human RGCs, which are difficult to obtain and maintain *in vitro*. Indeed, RGCs constitute less than 1% of all retinal cells in the adult eye [3,4], making the prospect of isolating large numbers of human RGCs from donor tissue challenging. Human pluripotent stem cells (hPSCs) including both embryonic stem cells (hESCs) and induced pluripotent stem cells (hiPSCs), represent a promising alternative source for generating human RGCs. For RGCs, two challenges remain: (i) generating sufficiently pure and high yield of RGCs, and (ii) maintaining them in culture in a manner that preserves their developmental trajectories and diverse subtypes. Previous studies identified 40-46 RGC subtypes in the mouse retina [5,6] and 18 subtypes in primates [7], although the exact number in humans remains unknown. This diversity is biologically important: RGC subtypes differ in their morphology, function, and potentially disease vulnerability [8]. Hence, heterogeneity must be captured to allow the reliable study of disease mechanisms and therapeutic responses.

Many RGC differentiation protocols rely on using small molecules to induce two-dimensional (2D) differentiation with positive selection based on a limited set of surface markers [9–12], yet such strategies may bias cultures toward particular molecular subtypes. Additionally, these cultures lack the three-dimensional (3D) organisation and signalling gradients that drive retinogenesis *in vivo*, hence miss the spatial cues required for physiologically appropriate specification and maturation. Furthermore, flow-cytometric marker-based enrichment is rarely complemented by transcriptome-level validation, leaving uncertainties about subtype composition and cellular heterogeneity. Retinal organoids offer an opportunity to overcome these limitations by providing an *in vivo*-like environment that supports the emergence of laminated retinal architecture and intrinsic patterning cues that shape early RGC identity. Further, retinal organoids recapitulate the developmental hierarchy of retinogenesis, in which RGCs are the first neuronal population to emerge [13]. This temporal advantage allows the generation of comparatively enriched RGC populations when organoids are harvested at early stages. Building on this principle, we applied an RGC-enriched approach to examine RGC differentiation and overall cellular composition in hPSC-derived retinal organoids. Organoids were dissociated at day 40, corresponding to peak expression of *POU4F* (*BRN3*) transcription factors [14], which are key regulators of RGC differentiation, survival and function [15–17], and replated for an additional 14 days in medium formulated to support RGC survival and maturation. We used flow cytometry and single-cell RNA sequencing (scRNA-seq) to assess enrichment efficiency, sample-to-sample variability, and the spectrum of retinal and off-target cell types present in the resulting cultures. Our findings demonstrate that protein marker-based assessment alone can overestimate RGC identity and highlight the necessity of single-cell transcriptomic validation for accurate evaluation of hPSC-derived RGC differentiation.

## Methodology

### Ethics

This work was approved by the office of Research Ethics and Integrity of the University of Melbourne (2026-32991-75214-7), as per the requirements of the NHMRC, in accordance with the Declarations of Helsinki.

### hPSC maintenance

HESC H9 (WiCell) and hiPSC WAB-0222 [18] lines were maintained on Matrigel-coated plates (Corning, #354230) in mTeSR™ Plus medium (STEMCELL Technologies, #100-0276). Medium was changed every other day, and cells were passaged weekly for routine maintenance.

### Retinal organoid differentiation

Retinal organoids were generated following a previously published protocol [14], with some modifications. On Day-1, hPSC cultures at 70% confluency were dissociated using ReLeSR (STEMCELL Technologies, #100-0484), and 4,000-5,000 cells were seeded into each well of a 96-well low adhesion U-bottom plate (Corning, #7007) in mTeSR™ Plus containing 20 µM Y-27632 (STEMCELL Technologies, #72304). Plates were centrifuged at 300 × g for 3 minutes to promote aggregate formation. On Day 0, each well received 50 µL mTeSR™ Plus and 50 µL Neural Induction Medium (NIM; DMEM/F12 [1:1; Thermo Fisher Scientific, #11320033] supplemented with 1% N2 [Thermo Fisher Scientific, #17502001], 1% MEM non-essential amino acids [Sigma, #M7145], 1× penicillin-streptomycin [Gibco, #15140122], and 2 µg/mL heparin [STEMCELL Technologies, #7980]), supplemented with 20 µM Y-27632. On Day 1, 40-45 aggregates were transferred directly into 10-centimeter polystyrene dishes (Corning, #430591) and fed with 6 mL fresh mTeSR™ Plus and 6 mL NIM. Medium was refreshed with NIM on Days 2 and 3. On Day 6, cultures were changed to NIM containing 50 ng/mL BMP4 (R&D systems, #314-BP-010). On Day 8, wells of a 6-well plate were pre-coated with 500 µL fetal bovine serum (FBS; Cytiva HyClone, #SH30084.03), and 20-30 aggregates were plated per well. On Day 9, half of the BMP4-containing medium was replaced with NIM to achieve a final concentration of 10% FBS, followed by half-medium changes on Days 12 and 15. On Day 16, organoids were lifted and transferred into 10-centimeter dishes containing Retinal Differentiation Medium (RDM; DMEM/F12 [3:1], 2% B27 supplement [Life Technologies, #17504044], 1% MEM non-essential amino acids, and 1× penicillin-streptomycin) containing 1% FBS. RDM was replaced on Days 18 and 20 using media supplemented with 3% and 5% FBS, respectively. On Day 22, organoids were transitioned to Advanced RDM (ARDM; DMEM/F12 [3:1], supplemented with 2% B27, 1% MEM non-essential amino acids, 1× penicillin-streptomycin, 1× GlutaMAX, 10% FBS, and 100 µM taurine). Half-medium changes were performed every 2-3 days until Day 40.

On Day 40, retinal organoids were dissociated using the Papain Dissociation Kit (Worthington Biochemical Corporation, #LK003153). Following a 30-minute incubation at 37 °C, organoids were triturated 20 times with a 1-mL pipette to generate a suspension enriched for small aggregates. The suspension was transferred to a 15-mL tube containing an equal volume of 10 mg/mL ovomucoid protease inhibitor and centrifuged at 300 × g for 5 minutes. The resulting pellet was resuspended in 1 mL Neurobasal-based neuronal differentiation medium (NDM; Neurobasal Plus [Thermo Fisher Scientific, #A3582901], supplemented with 1% MEM non-essential amino acids, 1% GlutaMAX [Thermo Fisher Scientific, #35050061], 1% 45% glucose [Merck, #G8769], 1× penicillin-streptomycin, 1× B27, 1× N2, 1× CultureOne [Thermo Fisher Scientific, #A3320201], and 1× Normocin [InvivoGen, #ant-nr-2]). All supplements were added fresh immediately before use. Viable cells were counted using trypan blue exclusion and plated onto 12-well plates coated with poly-D-lysine (2 µg/cm²; Sigma-Aldrich, #P0899-10MG) and laminin (1 µg/cm²; Sigma-Aldrich, #L2020-1MG) at a density of 200,000 cells/cm². Following dissociation, cells were cultured in NDM containing 20 µM Y-27632, 10 ng/mL CNTF (PeproTech; #450-13-50UG), 40 ng/mL BDNF (PeproTech, #450-02-50UG), 10 µM forskolin (Biogem/Lonza, #6652995), and 3 µM DAPT (Abcam, #AB120633), applied as a full medium change. Y-27632 was added immediately after plating. On Day 41, a half-medium change with NDM containing CNTF, BDNF, forskolin, and DAPT was performed, with medium added gently along the well perimeter to avoid dislodging adherent cells. A full medium change was performed on Day 43 to remove debris and eliminate residual Y-27632. From Days 44-47, half-medium changes were performed every 2-3 days while maintaining CNTF, BDNF, forskolin, DAPT, and CultureOne. From Days 48-54, medium changes continued as above, with forskolin increased to 25 µM; Day 48 involved a full medium change, followed by half-medium changes every 2-3 days.

### Flow cytometry

hPSC-RGCs cultured for 14-16 days on PDL/laminin-coated 24-well plates were dissociated using either TrypLE Express (Thermo Fisher Scientific; 12604-021, 10 minutes, 37 °C) or the Papain Dissociation Kit (Worthington Biochemical Corporation; LK003153, 20 minutes, 37 °C), depending on cell numbers. The papain kit was more effective at preserving cell viability in cultures with lower cell yields. Cells were collected by centrifugation at 340 × g for 5 minutes at 4 °C, and the resulting pellets were resuspended in DPBS-2% BSA (FACS buffer). Live/dead staining was performed using the Fixable Violet 405 dye kit (Thermo Fisher Scientific, #L34964, 30 minutes, room temperature). Cells were then washed in the FACS buffer, and centrifuged at 340 × g for 5 minutes at 4 °C. For extracellular staining, cells were resuspended in FACS buffer and incubated with Anti-CD90 BV510 (BD Horizon, #563070) for 45 minutes on ice. For intracellular staining, cells were first resuspended in DPBS, fixed with 4% paraformaldehyde for 10 minutes at room temperature, and permeabilized in 0.1% BSA in DPBS-0.05% Triton X-100-0.1% Tween-20 for 15 minutes at room temperature. Cells were subsequently resuspended in 0.1% BSA-DPBS-0.1% Tween-20 and incubated with the following antibodies for 45 minutes at room temperature: Anti-SNCG Alexa Fluor 488 (Santa Cruz, #sc-65979), Anti-ISL1 PE (BD Pharmingen, #562547), Anti-GFAP Alexa Fluor 647 (BD Pharmingen, #560298), and Anti-POU4F Alexa Fluor 647 (Santa Cruz, catalog #sc-390780). Antibody selection was based on markers that had been detected in hPSC-derived RGCs. The pan-RGC marker RBPMS was therefore not included due to its variable and often low expression in these cultures [19]. Stained cells were analysed using a CytoFLEX LX flow cytometer, and data were processed with FlowJo v10.10 software. A negative control using hPSC-derived RPE cells is shown in **Fig. S1**.

### Single cell preparation of iPSC-RGCs

hPSC-RGCs cultured for 14-16 days on PDL/laminin-coated 24-well plates were dissociated using the Papain Dissociation kit according to the manufacturer’s protocol. Briefly, cells were incubated with papain/DNase solution at 37 °C for 20 minutes, gently triturated, and the enzymatic reaction was quenched using the albumin-ovomucoid inhibitor solution. Following dissociation, cells were centrifuged and washed in 1% BSA, then sequentially filtered through 30 µm pre-separation filters (Miltenyi Biotec; 130-041-407) and kept on ice. Cell viability and concentration were determined by Trypan Blue exclusion using a Countess 3 FL Automated Cell Counter (Thermo Fisher; AMQAF2000). Pelleted cells were fixed for long-term storage using the GEM-X Flex Sample Preparation v2 Kit (10x Genomics; 1000781) according to the manufacturer’s CG000782 protocol for GEM-X Flex Gene Expression.

### Generation of single cell GEMs and sequencing libraries

Single-cell suspensions were processed by the Australian Genome Research Facility using the 10x Genomics Chromium GEM-X Flex Gene Expression Human 4-plex assay according to the manufacturer’s protocol. For each sample, 3 × 10L fixed cells were hybridised with uniquely barcoded probe sets (BC001-BC004) for 20 hours at 42 °C, washed, and pooled at equal concentrations. Pooled cells were loaded onto a Chromium X instrument with GEM-X FX Chips, combining barcoded Gel Beads, master mix, and Partitioning Oil B to generate single-cell Gel Beads-in-Emulsion (GEMs) targeting recovery of ∼80, 000 cells. GEMs were transferred to a thermal cycler to ligate the left-hand and right-hand probes that remained hybridised to their target RNA, hybridise Gel Bead primers to the capture sequence of the ligated probe pairs, and extend barcode sequences. Following emulsion breaking with Recovery Reagent, the ligated and extended products were PCR-amplified, cleaned, and indexed. Libraries were quality-assessed on a TapeStation D1000, quantified by qPCR, and sequenced on an Illumina NovaSeq X Plus (10B flow cell; 150 bp paired-end + 10 % PhiX).

### Mapping of reads to transcripts and cells

Base-call files were processed using Cell Ranger v7.1.0 (10x Genomics) configured for the Chromium Single Cell 3′ v3.1 chemistry. Reads were aligned to the *Homo sapiens* reference genome (GRCh38, Ensembl release 109). Cell Ranger performed default barcode and UMI correction to generate unfiltered gene-by-cell count matrices. No library aggregation was performed.

### scRNA-seq processing, integration, and visualisation

#### Cell recovery and ambient RNA correction

Raw gene-barcode count matrices were processed to remove empty droplets and correct ambient RNA contamination. Ambient RNA contamination was corrected using the *SoupX* package [20]. For each sample, the ambient RNA profile was estimated from barcodes not present in the cell-containing matrix (raw-only barcodes) with total UMI counts in the range 1-100 UMIs. Non-empty putative cells were pre-clustered in *Seurat* (v5) using LogNormalize → FindVariableFeatures → ScaleData → Run principal component analysis (PCA; 30 PCs) → FindNeighbours → FindClusters (Leiden, resolution = 0.4). These quick cluster labels were supplied to *SoupX*, contamination fraction (ρ) was estimated using *autoEstCont*, and corrected counts were generated with adjustCounts.

#### Post-processing and doublet detection

Based on the pre-filtering quality-control metrics (**Fig. S2**), *Seurat* objects were reconstructed from SoupX-corrected counts and filtered using thresholds of nFeature_RNA 1000-8000 and mitochondrial RNA percentage < 30%. Doublets were detected on raw counts using *scDblFinder* (SingleCellExperiment backend; serial execution) [21], and only singlets were retained.

#### Normalisation and Harmony Integration

Each sample was normalised independently using SCTransform v2 (glmGamPoi backend) [22], regressing out mitochondrial transcript fraction. After merging samples, principal component analysis was performed on SCT residual features, and sample-associated technical variation was mitigated using Harmony. Harmony aligns transcriptionally similar cell states across samples without assuming technical replicates or enforcing alignment of non-overlapping populations. Harmony embeddings were used for UMAP visualisation and construction of the shared nearest-neighbour graph (dims 1-30). Clustering was performed using the Leiden algorithm across a range of resolutions (0.1-1.0) to assess cluster stability and granularity.

#### Dimensionality reduction and clustering

Principal component analysis (PCA; 50 components) was performed on the Harmony-integrated object. To determine the number of biologically informative dimensions, we primarily examined the Elbow plot, which revealed a clear inflection point at approximately 20-25 principal components (**Fig. S3A**). This observation was supported by the cumulative variance plot, which showed that the majority of structured variation was captured within the first ∼20-25 PCs, with progressively smaller gains thereafter (**Fig. S3B**). To ensure retention of potentially informative higher-order components while avoiding excessive noise, a conservative cutoff of the first 30 PCs was selected for all subsequent neighbour graph construction, UMAP embedding, and clustering.

#### Cluster stability assessment and resolution selection

UMAP embeddings were generated using Harmony-corrected embeddings (min.dist = 0.3, spread = 1.0) with fixed random seeds to ensure reproducibility [23]. Graph-based clustering was performed using the Leiden algorithm (algorithm = 4) across a range of resolutions from 0.1 to 1.0 in increments of 0.1 [24]. Cluster stability across resolutions was assessed by calculating the Adjusted Rand Index (ARI) between clustering solutions at consecutive resolutions. ARI values increased rapidly between low resolutions (0.2-0.4) and reached a high, stable range from resolution 0.4 onwards, indicating robust preservation of cluster structure across increasing granularity (**Fig. S4A**). Consistent with this, cluster visualisation demonstrated coherent cluster propagation with minimal fragmentation across intermediate resolutions, forming a stable plateau between resolutions 0.4 and 0.5 (**Fig. S4B**). Based on the convergence of ARI stability and clustree topology, a resolution of 0.5 was selected for all downstream analyses.

#### Differential gene expression and marker identification

Differential expression analysis was performed on the Harmony-integrated object using Seurat v5. Prior to differential expression analysis, clusters containing fewer than 100 cells were excluded to ensure sufficient statistical power. The object was prepared for DE using PrepSCTFindMarkers(). Cluster-specific markers were identified using FindAllMarkers (test.use = “wilcox”, only.pos = TRUE, min.pct = 0.10, log2FC threshold = 0.25), comparing each cluster against all other cells. Gene identifiers were standardised to HGNC symbols using Ensembl v109 via biomaRt.

#### Cell type annotation and marker validation

Gene symbols were standardised to HGNC conventions using HGNChelper [25] and biomaRt [26]. For each Seurat cluster, average gene expression (SCT assay) was z-scored and visualised with ComplexHeatmap to confirm marker specificity and to guide manual annotation. Clusters containing fewer than 100 cells were excluded. The remaining clusters were assigned to retinal progenitor cells, RGCs, amacrine cells, horizontal cells, photoreceptors, retinal pigment epithelium (RPE), Multilineage (stressed), Other (*HOX*-enriched), or Other based on the expression of established retinal marker genes from human datasets (Table 1) [27–29].

**Table 1:** Established retinal marker genes from human datasets.

| Cell Identity | Marker genes |
| --- | --- |
| Retinal progenitor cell | *SOX2, PAX6, VSX2, LHX2, MKI67, ATOH7, FABP7, HES6, DLL3, PTF1A* |
| RGC | *POU4F2 (BRN3B), ISL1/2, EOMES, RBPMS, SNCG, THY1, NEFL, NEFM, GAP43, SLC17A6, POU6F2, ELAVL3, ELAVL4* |
| Amacrine cell | *GAD1, TFAP2A, PRDM13, CHAT, SLC6A9* |
| Horizontal cell | *PROX1, LHX1, ONECUT1, ONECUT2, GJA10, RORB* |
| Photoreceptor | *OTX2, CRX, THRB, NRL, RHO, GNAT1, NR2E3, ARR3, GNAT2, PDE6H* |
| RPE | *RPE65, MITF, TTR, TYR, TYRP1, RDH5* |
| Multilineage (stressed) | *FOS, JUN, ATF3, DDIT3, CDKN1A, HSPA1B* |
| Other (*HOX*-enriched) | *HOXB5-8* |

### Projection of Multilineage (stressed) cells

To infer the lineage affiliation of “Multilineage (stressed)” cells, we applied Seurat’s label-transfer framework using non-stressed cells as the reference population. Cells in cluster 19, defined based on Harmony-integrated clustering, were treated as the query set. Transfer anchors were computed using PCA-based reduction on normalised expression profiles, and lineage predictions for each stressed cell were generated using TransferData.

#### Trajectory inference, lineage reconstruction, and pseudotime estimation

The developmental root was defined by proliferative activity within retinal progenitor cell clusters. Cell-cycle phase was assigned using Seurat’s CellCycleScoring with canonical S-phase and G2/M gene sets. Clusters 15 and 17 exhibited high proliferative activity; however, cell-cycle phase analysis revealed distinct cycling states. Cluster 15 comprised a mixed cycling population (50.9% S phase, 49.1% G2/M), whereas cluster 17 was almost exclusively G2/M (98.9%), indicating a highly synchronised mitotic progenitor population. Cluster 17 was therefore selected as the trajectory root, as it represents the most actively cycling progenitor state. In contrast, differentiated neuronal populations, including RGCs (61-95.8% G1), interneurons (75.8-84.6% G1), and photoreceptor-committed cells (53.8-71.2% G1), were predominantly in G1, consistent with post-mitotic states.

Cell differentiation trajectories were inferred using *Slingshot* (v2.16.0) applied to the PCA embeddings derived from the Harmony-integrated object (R v4.5.1) [30]. Trajectory inference was performed in a family-wise manner, with separate analyses conducted for major retinal lineages without cross-lineage interference. For each lineage family, Slingshot inferred smooth principal curves through the cluster topology, generating continuous pseudotime values for all included cells. Pseudotime distributions for downstream comparison and visualisation were summarised using ridge plots (ggridges).

#### RGC Subclustering and Subtype Annotation

Clusters 2, 3, 11, and 14 were identified as RGCs and extracted for downstream analysis. All analyses were performed in R (v4.5.1) using Seurat (v5) together with *harmony*, *mclust*, *clustree*, *dplyr*, and *ggplot2*. This RGC population was reprocessed independently. The RNA assay was re-normalised using SCTransform, and PCA was performed on the resulting SCT assay, computing 50 PCs. Dimensionality selection was guided by PCA elbow plots as well as variance explained and cumulative variance plots (**Fig. S5**). Based on these metrics, the first 20 PCs were retained for downstream neighbour graph construction and dimensionality reduction. A shared nearest-neighbour graph was constructed using the Harmony embeddings (dimensions 1-20), and Leiden clustering was applied across a resolution grid ranging from 0.1 to 1.0 (step size = 0.1). Clustering stability was assessed by calculating the ARI between adjacent resolutions using *mclust* and by visual inspection of cluster relationships using clustree. Based on ARI profiles and clustree topology, a final clustering resolution of 0.3 was selected (**Fig. S6**).

RGC subtype markers were selected based on previously reported human RGC marker genes associated with major RGC subtypes (Table 2), including α/Parasol, Midget, direction-selective ganglion cells (DSGC), orientation-selective ganglion cells (OSGC; J-RGC-like), large sparse RGCs, and intrinsically photosensitive RGCs (ipRGCs) [7,29,31–33].

**Table 2:**
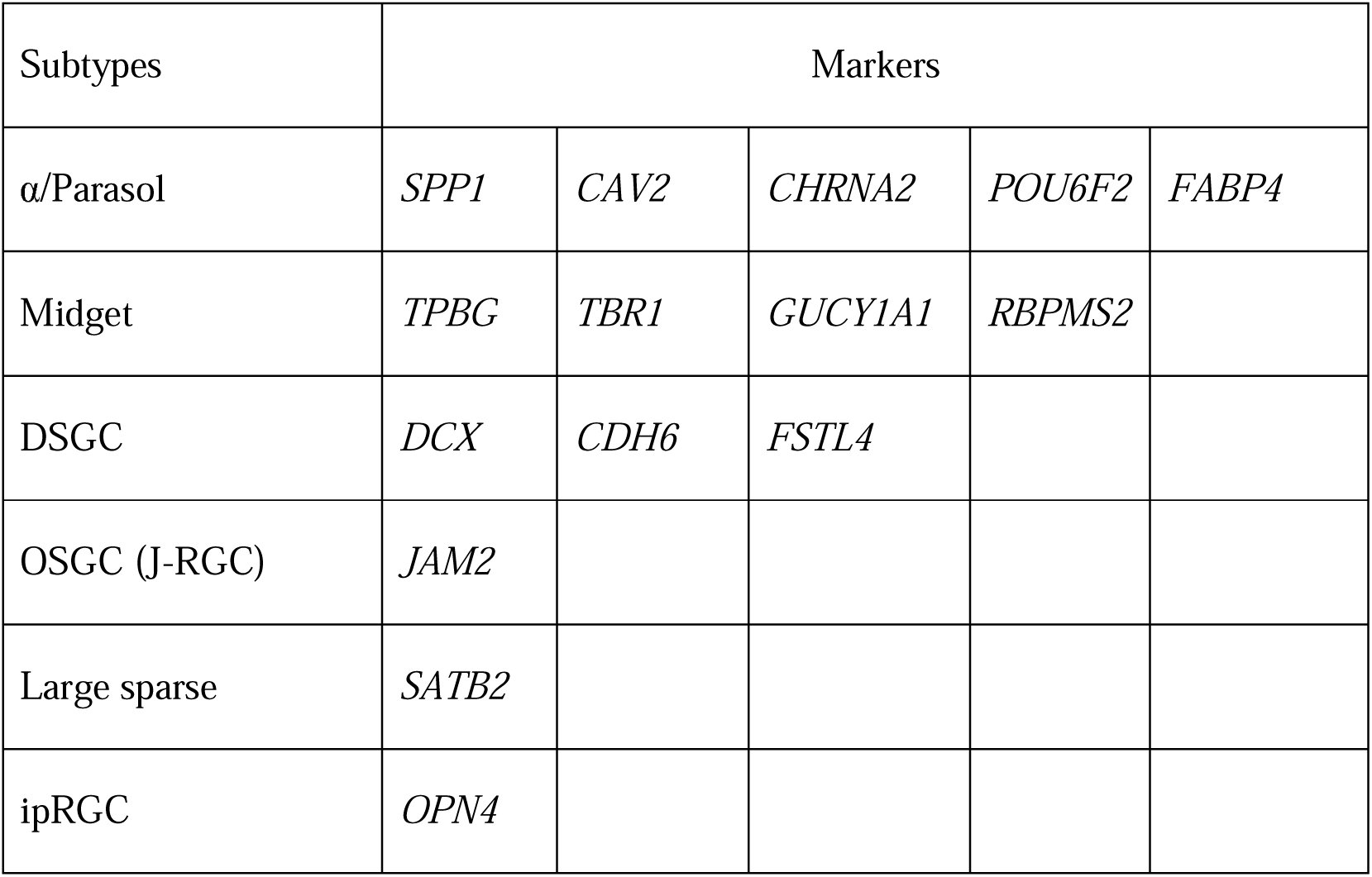
Established marker genes for human RGC subtypes.

## Results

### Generation of RO and incorporation of RGC enrichment strategies

The RGC enrichment approach was developed by integrating elements from prior studies (**Fig. 1A,B**). Retinal organoids were first generated using a modified version of a previously established protocol [14], then dissociated and plated for further differentiation, a step that has been shown to favour enrichment of neuronal populations expressing CD90 [10]. Retinal organoids were dissociated at day 40, a developmental stage reported to exhibit peak *POU4F* expression, a canonical RGC marker [14]. Finally, cultures were maintained under conditions previously shown to support the survival and maturation of emerging RGC-like cells [9](**Fig. 1B**).

**Figure 1.**
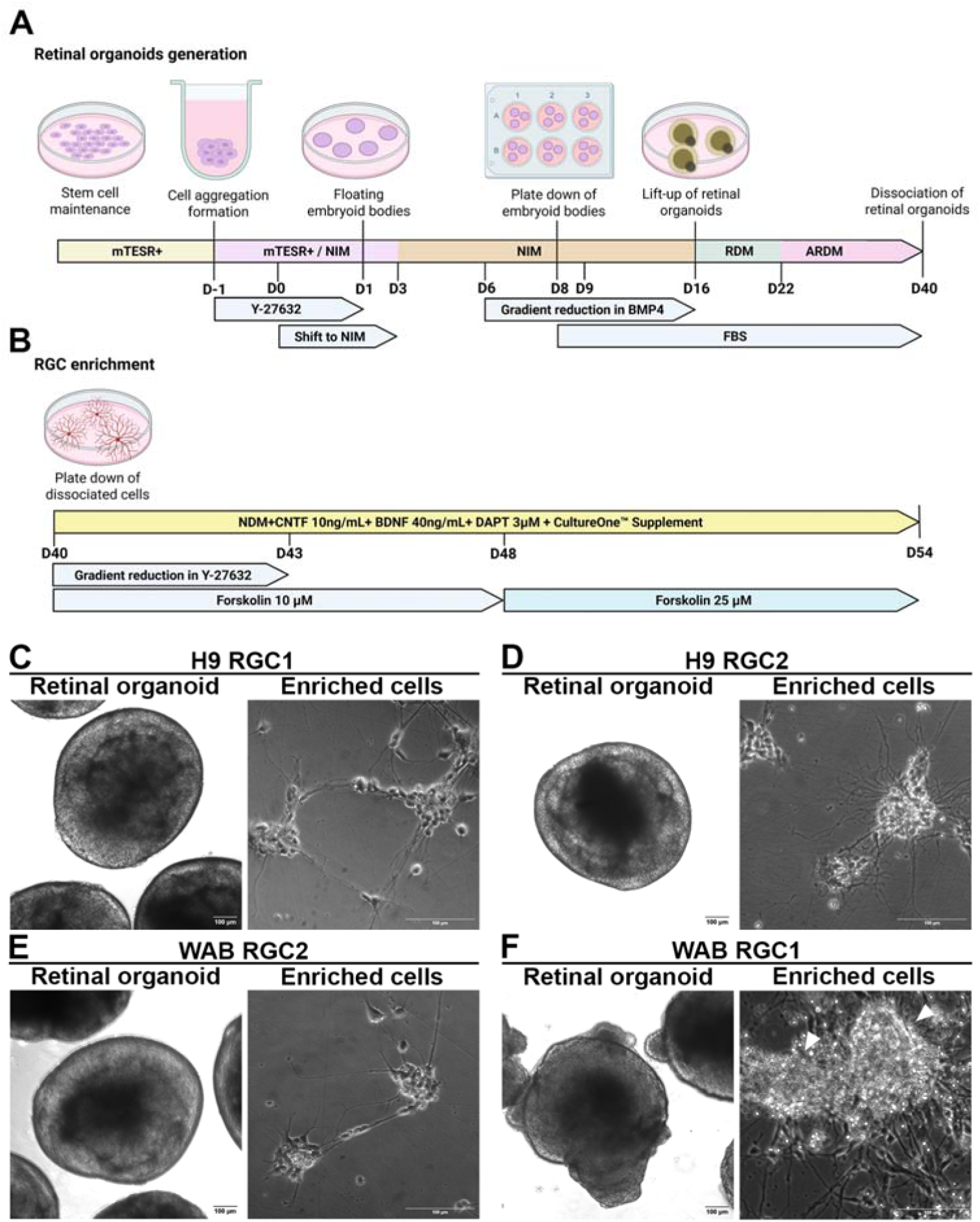
Generation of retinal organoids and subsequent RGC enrichment. (**A**) Schematic diagram illustrating the generation of retinal organoids, and (**B**) the subsequent enrichment of RGCs following dissociation of retinal organoids at day 40 (D40). Created with BioRender.com (**C-F**) Brightfield images showing retinal organoids at D40 (left panel) and the corresponding RGC-enriched enriched cell populations at D54 (right panel) for (**C**) H9_RGC1, (**D**) H9_RGC2, (**E**) WAB_RGC2, (**F**) WAB_RGC1. Scale bar: 100 µm.

Retinal organoids derived from two hPSC lines were produced in duplicate independent cultures (H9_RGC1, H9_RGC2, WAB_RGC1, and WAB_RGC2). All samples underwent the same differentiation procedure from the hPSC stage to retinal organoids formation. Typically, lifted retinal organoids after day 16 acquire a smooth, rounded morphology with a distinct bright halo along the periphery, indicating the establishment of laminated retinal layers. This characteristic morphology was observed in H9_RGC1, H9_RGC2, and WAB_RGC2 (**Fig. 1C-E**, left panels). In contrast, WAB_RGC1 organoids became flattened during the plating-down period, likely as a result of disruption to the embryoid body surface during manual transfer. Despite the flattening, these organoids were lifted and retained for downstream culture and analysis to assess whether such morphological changes could serve as a potential morphological selection criterion in future protocols. Upon lifting, WAB_RGC1 organoids developed irregular, clumped edges around a central dense core, and lacked the characteristic bright peripheral halo, suggesting disrupted lamination (**Fig. 1F**, left panel). In addition, regions of folded or thickened tissue were observed on the surface of WAB_RGC1 organoids, indicating a failure to establish proper neuroepithelial polarity (**Fig. 1F**, left panel). We therefore included WAB_RGC1 as a suboptimal differentiated control to assess whether altered organoid morphology would influence RGC yield.

After 14 days of RGC enrichment following organoid dissociation, cells from H9_RGC1, H9_RGC2, and WAB_RGC2 formed multiple well-defined neuronal clusters interconnected by thin, long dendritic processes, typical of neuronal networks (**Fig. 1C-E**, right panels). By contrast, WAB_RGC1-derived cultures displayed multiple dense aggregates containing several large soma-like centres with multiple extended multiple neurites (**Fig. 1F**, right panel; large soma-like centres indicated by arrowheads).

### Flow cytometric assessment of RGC marker expression

To quantify the abundance of RGC-like cells within each enriched culture, flow cytometry analysis was performed using four markers, which in combination are indicative of RGC identity: POU4F (recognising POU4F family proteins), ISL1, SNCG, and THY1 (**Fig. 2A; Fig. S7**). All samples demonstrated high proportions of POU4F+ cells (79.0 to 95.1%), consistent with efficient induction of sensory neurons/ RGC lineage identity. SNCG, a small cytosolic protein highly enriched in RGCs [34], was used as an additional marker of RGC differentiation. ISL1, a transcription factor expressed in RGCs and other inner retinal neurons [35,36], showed greater variability across samples (18.0-58.0%), reflecting the known stage-dependent temporal dynamic expression of ISL1 during RGC differentiation [35]. SNCG expression was similarly high in H9_RGC1, H9_RGC2 and WAB_RGC2 (81.3-91.0%), but was markedly reduced in WAB_RGC1 (21.5%), consistent with its less organised neurite architecture. THY1, a surface protein historically used as an RGC antigen in the retina [37], displayed the greatest variability, ranging from 3.7% in WAB_RGC2 to 29.3% in H9_RGC1. This variability aligns with the recognised instability of THY1 as a surface marker as it labels only a subset of RGCs and is also detectable in non-RGC retinal cell types [38]. Together, these data indicate the presence of RGC-associated marker expression across all samples, with some variability among individual markers.

**Figure 2.**
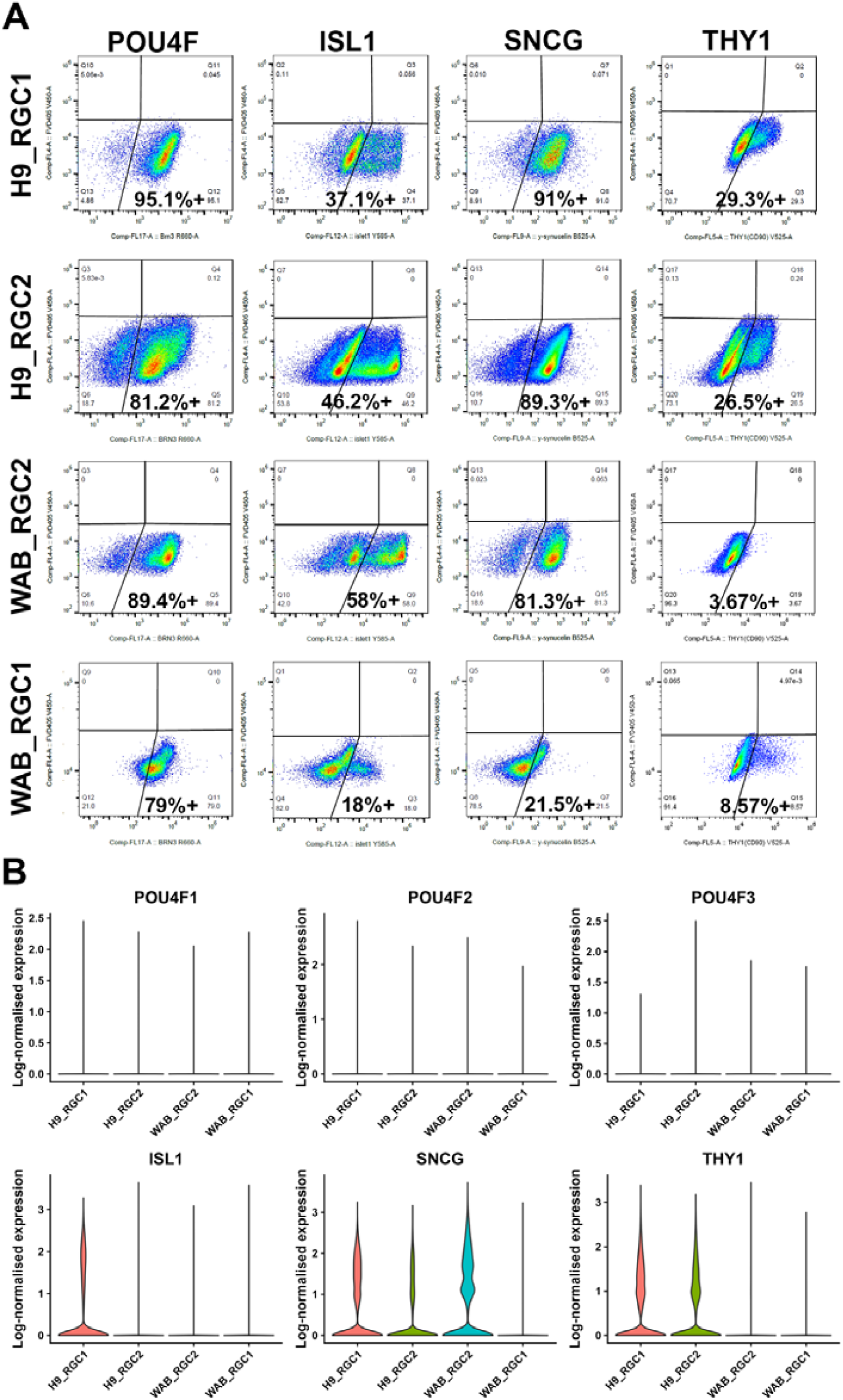
Flow cytometry assessment of RGC-associated marker expression. **(A)** Representative flow cytometry dot plots showing expression of POU4F, ISL1, SNCG, and THY1 across four samples (H9_RGC1, H9_RGC2, WAB_RGC2, and WAB_RGC1). Rows correspond to individual samples and columns to individual markers. Gates were defined using unstained populations. Percentages denote the fraction of marker-positive cells within the live singlet population. **(B)** Violin plots showing log-normalised mRNA expression of canonical RGC markers *POU4F* family members (*POU4F1*, *POU4F2*, *POU4F3*), *ISL1*, *SNCG,* and *THY1* across H9_RGC1, H9_RGC2, WAB_RGC2, WAB_RGC1. Each violin represents the distribution of transcript abundance across individual cells.

All RGC markers used for flow-cytometry validation were also detected at the transcript level in the scRNA-seq dataset; however, the mRNA expression of these individual markers varied substantially across all samples (**Fig. 2B; Fig. S8**). *POU4F1*, *POU4F2*, and *POU4F3* transcripts were detected in only a small fraction of cells across all samples, with the majority of cells exhibiting zero or near-zero expression. *ISL1* mRNA was detected in a small subset of cells primarily in H9_RGC1, while H9_RGC2, WAB_RGC2, and WAB_RGC1 showed little to no detectable expression. *SNCG* transcripts were detected in subsets of cells in H9_RGC1, H9_RGC2, and WAB_RGC2, with a distinct population exhibiting elevated expression, whereas *SNCG* expression was minimal in WAB_RGC1, where most cells showed near-zero transcript levels. *THY1* transcripts were present in subsets of cells in H9_RGC1 and H9_RGC2 but were largely absent in WAB_RGC samples, with only rare cells showing detectable expression. These findings indicate that reliance on flow cytometry, through detection of stable or residual protein, may overestimate RGC abundance relative to scRNA-seq.

### Identification and characterisation of 22 subpopulations

Across the four samples, a total of 73,642 high-quality single cells were retained for downstream integrated analysis (**Table S1**). At a clustering resolution of 0.5, a total of 22 transcriptionally distinct clusters were identified (**Fig. 3A**). However, cluster 22 contained fewer than 100 cells (n = 91) and was therefore excluded from downstream analyses. Based on marker expression, clusters were annotated into different cell types (**Fig. 3B**). Sample-level comparisons revealed notable differences in differentiation bias across the four datasets (**Fig. 3C**). In H9_RGC1, RGCs constituted the largest population (45%), followed by amacrine cells (15%), retinal progenitor cells (14%), *HOX*-enriched cells (13%), and horizontal cells (7%). In H9_RGC2, *HOX*-enriched cells were the most abundant (30%), with RGCs and amacrine cells each accounting for 19%, followed by retinal progenitor cells (11%), and photoreceptor-committed cells (10%). WAB_RGC2 displayed 38% of cells classified as photoreceptor-committed cells and 27% as RGCs, alongside horizontal cells (12%), other minor populations (8%), retinal progenitor cells (7%), and amacrine cells (6%). WAB_RGC1 exhibited the highest proportion of retinal progenitor cells (47%), with RGCs (10%), amacrine cells (10%), RPE (9%), photoreceptor-committed cells (7%), *HOX*-enriched cluster (5%), and other populations (5%) (**Table S2**).

**Figure 3.**
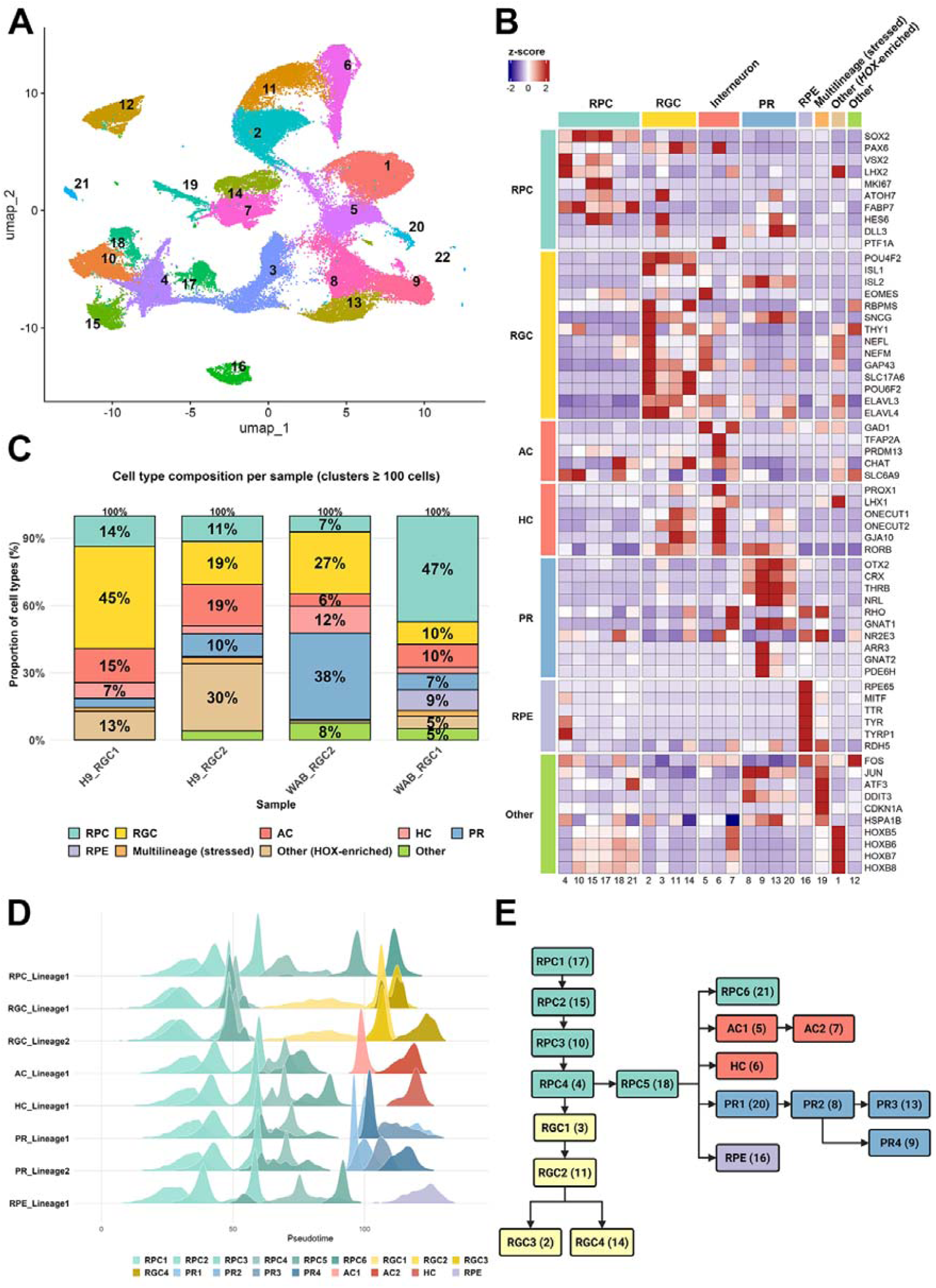
Single-cell transcriptomic profiling of RGC-enriched dissociated retinal organoid cells. (**A**) UMAP visualisation showing the distribution of the 22 transcriptionally defined clusters identified in the integrated dataset. (**B**) Heatmap displaying z-scored average log-normalised RNA expression of different marker genes across clusters. Rows correspond to marker genes grouped by major retinal cell classes, and columns represent the 22 clusters. Left annotation indicates the retinal cell type associated with each marker set, and the top annotation reflects the assigned cell-type identity for each cluster. (**C**) Stacked bar charts showing the proportional composition of annotated retinal cell types in each sample. (**D**) Ridge plots illustrating pseudotime distributions for the major developmental lineages. Each ridge reflects the relative density of cells and the progression of cells along pseudotime within a given lineage. (**E**) Lineage tree reconstructed from pseudotime analysis, showing hierarchical branching of retinal progenitor populations into downstream neuronal and epithelial fates. Each node is labelled with its cluster identifier, with bracketed numbers indicating the corresponding cluster IDs. RPC: retinal progenitor cell, RGC: retinal ganglion cell, AC: amacrine cell, HC: horizontal cell, PR: photoreceptor, RPE: retinal pigment epithelium.

Six retinal progenitor cells clusters (4, 10, 15, 17, 18, 21) were annotated based on the expression of canonical retinal progenitor cell markers such as *VSX2*, along with key transcription factors regulating neuronal fate, including *PAX6* and *SOX2 [39–41].* They also expressed *LHX2*, which is essential for maintaining open chromatin during retinogenesis and for gliogenesis [42,43]. Cell cycle-related genes were unevenly distributed among progenitor subpopulations, with the G2/M phase marker *MKI67* showing predominant expression in clusters 15 and 17.

RGC clusters were annotated based on the expression of canonical markers including *POU4F2*, *ISL1*, *ISL2*, *EOMES*, *SNCG*, *THY1*, *NEFL*, *NEFM*, and *RBPMS* [5,44–47]. Using these markers, four transcriptionally distinct RGC subpopulations were identified (**Fig. 3B**), All of these clusters (2, 3, 11, and 14) expressed *POU4F2*. Cluster 3 represented the earliest RGC-fated state, characterised by high expression of *ATOH7*, *HES6*, and *DLL3*. Clusters 2 and 14 showed strong co-expression of *POU4F2*, *ISL1*, *RBPMS*, *NEFL*, *NEFM*, *SNCG*, *THY1*, *SLC17A6*, and *GAP43*, indicative of maturing RGCs. Clusters 11 and 14 exhibited partial expression of horizontal cell-associated markers such as *ONECUT1* and *ONECUT2*, indicating transcriptional heterogeneity within the enriched RGC population.

Two amacrine cell clusters (5 and 7) were enriched with an inhibitory neuron marker *GAD1* [27], while a horizontal cell cluster (6) expressed *PROX1*, *ONECUT1*, *ONECUT2* and *GJA10* [48–50]. Photoreceptor-committed cells identity was defined using established lineage markers, including *OTX2*, *CRX*, *THRB,* and *NRL*, together with rod-specific markers (*RHO*, *GNAT1*, *NR2E3*) and cone-specific markers (*ARR3*, *GNAT2*, *PDE6H*) [51–55]. Four clusters (8, 9, 13 and 20) showed strong expression of *OTX2* and *CRX*, confirming photoreceptor lineage commitment. Among these, cluster 9 displayed the most advanced phototransduction programme, co-expressing cone-specific markers (*ARR3*, *GNAT2*, *PDE6H*) together with rod-associated genes (*NRL*, *PDE6B*, *GNAT1*), indicating a population of mixed rod-cone photoreceptor precursors. Additionally, cluster 16 showed the expression of RPE*65*, *MITF, TYR*, *TYR* and *RDH5*, classical markers of RPE [56].

Cluster 19 exhibited a stress-associated transcriptional profile characterised by high expression of *FOS*, *JUN*, *ATF3*, *DDIT3*, *CDKN1A*, and *HSPA1B [57–61]*. This cluster was annotated as a Multilineage (stressed) population, and it was less than 5% of the total cells across all samples respectively. Differential expression analysis revealed induction of genes associated with apoptosis initiation, cellular stress signalling and early cell-death pathways, although no coherent set of canonical lineage markers was detected. To determine the underlying lineage composition masked by stress-induced transcriptional reprogramming, label-transfer analysis was performed (**Table 4**). This analysis revealed that cluster 19 comprised mainly of retinal progenitor cells, followed by “Other” cell types, RPE, *HOX*-enriched cells, and amacrine cells (**Table 4**).

**Table 4:** Lineage composition of the Multilineage (stressed) cluster identified by label-transfer analysis.

| **Family** | **# of cells** | **Percentage of cells (%)** |
| --- | --- | --- |
| **Retinal progenitor cell** | 483 | 35.3 |
| **RGC** | 43 | 3.1 |
| **Amacrine cell** | 125 | 9.1 |
| **Horizontal cell** | 14 | 1.0 |
| **Photoreceptor-committed cell** | 28 | 2.0 |
| **RPE** | 144 | 10.5 |
| **Other (*HOX*-enriched)** | 136 | 9.9 |
| **Other** | 397 | 29.0 |

Cluster 1 was annotated as *HOX*-gene-enriched cells, marked by high expression of *HOX*-related genes such as *HOXB5-8*. These cells also expressed neuronal structural genes such as *NEFL*, *NEFM* and *ELAVL4* but lacked retinal identity markers, indicating the formation of off-target posterior neural cell types not belonging to anterior retinal tissue. *HOX*-enriched populations were most abundant in the two H9-derived samples. In addition, cluster 12 was annotated as “Other” as it did not exhibit retinal cell markers.

### Pseudotime analysis reveals progressive retinal lineage trajectories

To reconstruct differentiation pathways across retinal lineages, we performed pseudotime and lineage trajectory analysis on the Harmony-integrated dataset comprising all four samples. Clusters annotated as “Multilineage (stressed)” and “Other” were excluded as they exhibited mixed lineage marker expression and/or off-target transcriptional programs, rather than a single coherent retinal identity. Inclusion of these cells could distort transcriptional similarity relationships and lead to artificial trajectory splits that do not reflect genuine developmental lineage decisions. Cells were ordered along developmental trajectories based on transcriptional similarity. Ten major lineages were identified, corresponding to retinal progenitor, RGC, amacrine, horizontal, photoreceptor, and RPE fates, each branching from the retinal progenitor cell cluster 17, which showed the highest expression of the proliferation marker *MKI67* (**Fig. 3D**). The earliest pseudotime positions were occupied by the RPC1-RPC3 populations, which together formed the central developmental trunk. RPC4 predominantly gave rise to two RGC lineages and to RPC5, which forms the root of all other major retinal cell lineages, including an additional retinal progenitor cell lineage, an amacrine cell lineage, one horizontal lineage, two photoreceptor lineages, and one RPE lineage (**Fig. 3E**).

### Transcriptional characterisation of RGC subtypes

To further resolve the molecular diversity within the RGC lineage, we re-clustered cells from RGC-associated clusters (2, 3, 11 and 14) at a resolution of 0.3, resulting in seven transcriptionally distinct RGC subclusters (**Fig. 4A**, **B**). Canonical human RGC markers [7,29,31–33] (**Table 2**) were detected across these subclusters, including markers for parasol, midget, DSGC, OSGC, and large sparse RGCs (**Fig. 4C**), suggesting that the cultures generate multiple RGC subtypes. Most subclusters showed overlapping expression of several RGC subtype marker genes, suggesting incomplete transcriptional segregation. Although clusters 1 and 4 showed elevated *OPN4* signal (**Fig. 4C**), it was detected in only 49 cells for cluster 1, and 11 cells in cluster 4 in the log-normalised RNA matrix. As this proportion fell below the minimum detection threshold (≥5% of cells expressing the marker), *OPN4* was not identified in the differential expression list (**Fig. 4C, D**). For each selected marker gene, cells with normalised expression greater than zero were considered positive, and Harmony-derived UMAP embeddings were used to visualise the spatial distribution of marker-positive cells (**Fig. 4D**).

**Figure 4.**
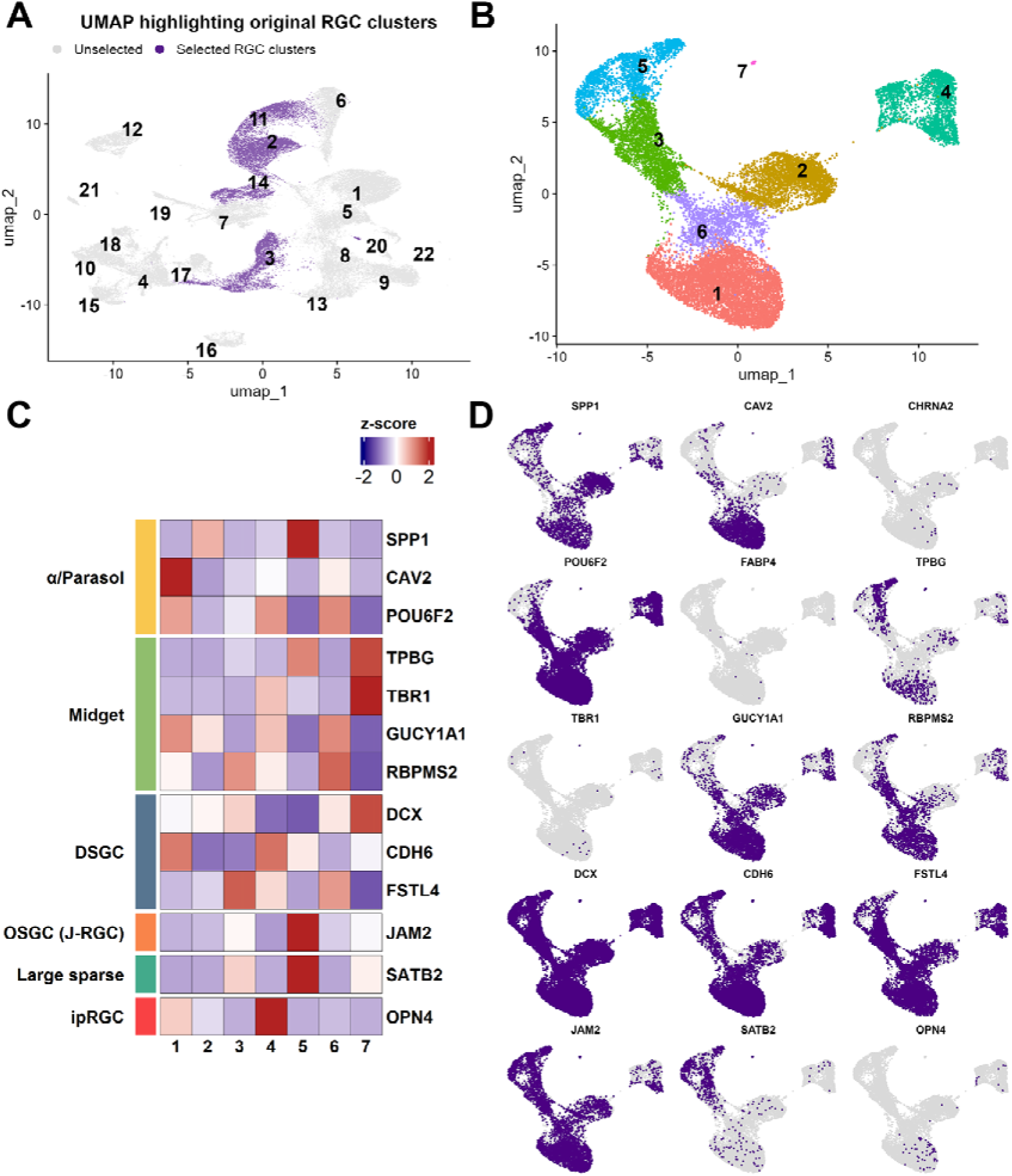
Subclustering and molecular characterisation of RGC subpopulations. (**A**) UMAP showing the clusters (in purple) selected for subsequent reclustering of the RGC subpopulation. (**B**) UMAP showing the seven distinct clusters identified. (**C**) Heatmap showing z-scored average log-normalised RNA expression of RGC subtypes markers. Rows represent marker genes grouped by major RGC subtypes, and columns represent the seven clusters. The left annotation indicates the subtype associated with each marker set. (**D**) UMAP visualisation of RGC subtype marker expression using log-normalised RNA expression values. For each marker, cells with detectable expression (log-normalised expression > 0) are highlighted in purple, while non-expressing cells are shown in grey. Each panel represents a single marker gene, with gene symbols displayed at the top of each plot.

Given that *OPN4* marks a rare subtype of RGC, we assessed whether additional rare melanopsin-expressing cells were missed due to normalisation and subsequent subclustering. Examination of the unnormalised, SoupX-adjusted RNA count matrix, where low-abundance transcripts remain easier to detect than in SCTransform-normalised data [62], identified 128 *OPN4*+ cells, comprising 120 cells with low transcript abundance and 8 cells with higher abundance (**Fig. S9 and Table S3**). Mapping these *OPN4*+ cells onto the Harmony-derived UMAP embedding revealed that approximately half were located within the horizontal cluster (cluster 6 of the full integrated RGC-enriched retinal organoid dataset; **Fig. 3**), while 51 cells were located within the RGC cluster (cluster 2; **Fig. 3**).

## Discussion

This work provides an in-depth analysis of RGC differentiation within hPSC-derived retinal organoid cultures using complementary flow cytometry and single-cell transcriptomic approaches. Discrepancies between protein- and transcript-based readouts highlight the limitations of marker-based assessments and underscore the importance of transcriptomic benchmarks for defining RGC identity in heterogeneous cultures. Across samples, protein-based assays suggested robust RGC enrichment, whereas transcriptomic profiling revealed a smaller RGC population alongside significant contributions from non-RGC neuronal populations. For instance, over 80% of cells were deemed POU4F+ and SNCG+ in H9_RGC2, yet scRNA-seq identified more than one-third of cells belonging to *HOX*-enriched cell clusters. These *HOX*-enriched clusters may represent posterior CNS-like neurons, such as spinal interneurons or hindbrain sensory populations, which are known to transiently express POU4F family proteins during development [63]. Similarly, WAB_RGC2 showed high proportions of POU4F+ cells and SNCG+ cells by flow cytometry, whilst scRNA-seq showed that the major cell types within this differentiation were from the photoreceptor lineage. These can also transiently express SNCG [64], which is widely expressed across multiple peripheral and central nervous system neuronal populations [65].

ISL1 expression further illustrates the limitations of protein-based classification, as it varies with cellular maturation and is not restricted to RGCs, being detectable in other neuronal populations [66]. This likely contributes to the variable proportion of ISL1+ cells observed across samples. In contrast, THY1+ cells consistently represented the smallest fraction, reflecting the dynamic nature of THY1 expression, its downregulation in stressed RGCs [67], and its presence in subsets of amacrine cells [68]. In H9_RGC1 and WAB_RGC2, scRNA-seq identified a greater proportion of RGCs than suggested by flow cytometry, highlighting the limited reliability of THY1 as a commonly used standalone marker for estimating RGC content [10]. These findings indicate that RGC-associated proteins can be detected outside the canonical RGC lineage, whereas scRNA-seq more accurately excludes non-RGC cells. This discrepancy reflects the broad and often transient expression of commonly used RGC markers during early neurogenesis, together with their persistence in off-target neuronal lineages. As a result, protein marker positivity alone can overestimate RGC identity when lineage resolution is incomplete. Future flow cytometry approaches could improve specificity by combining POU4F and SNCG with exclusion markers for photoreceptors and/or GAD1 for amacrine cells, thereby ensuring that marker-positive populations more faithfully represent RGCs.

Methodological differences between flow cytometry and scRNA-seq further contribute to these contrasting readouts. Flow cytometry enriches larger and healthier dissociated cells through gating and is highly sensitive to stable or residual protein, potentially enriching early neurons that transiently express RGC-associated markers, as discussed above. In contrast, scRNA-seq captures a broader and less selectively filtered population, including fragile or transcriptionally immature cells, and classifies identity based on integrated gene expression programs rather than individual markers. Although transcript dropout is an inherent limitation of scRNA-seq with typical protocols capturing only ∼10-20% of transcripts [69–71], reliance on multi-gene signatures reduces the impact of false negatives [72] and provides a more conservative and lineage-resolved definition of RGC identity.

Single-cell analysis resolved multiple RGC transcriptional states, alongside relatively higher proportions of retinal progenitor cells, off-target posterior neural populations, and photoreceptor-committed cells. Rather than forming a uniform population, RGCs in enriched cultures occupied a continuum of transcriptional states. Pseudotime reconstruction supported the coexistence of early, intermediate, and maturing RGC populations. Multiple RGC subtypes were also identified. Rare subtypes, including melanopsin-expressing RGCs, were detected at low abundance and were localised to specific transcriptional clusters, consistent with genuine low-level expression. However, these populations were particularly sensitive to analytical thresholds and normalisation strategies, underscoring the limitations of single-cell approaches for resolving rare neuronal populations and highlighting the importance of cautious interpretation when assessing low-abundance transcripts in developing systems.

In the presence of multiple developmental states within the culture, the observed overlap between RGC transcriptional programs and markers associated with other early-born retinal neurons can be understood in the context of retinal development. RGCs, amacrine and horizontal cells arise from shared *ATOH7*+ progenitors, and fate commitment during early retinogenesis occurs gradually rather than through abrupt transitions [73]. Transitional cells may therefore transiently activate transcriptional programs associated with multiple retina lineages before terminal differentiation. Consistent with this, developing RGCs or RGC-like neurons have been found to express markers associated with amacine and horizontal cells [74,75]. Hence, when interpreting the scRNAseq analysis, low-level or partial expression of lineage-associated markers should not be interpreted as definitive evidence of fate switching, but rather as a reflection of developmental immaturity or incomplete fate resolution.

This developmental heterogeneity complicates marker-based assessments of RGC enrichment. As enriched cultures contain early RGCs, maturing neurons, and diverse subtype identities, no single surface or intracellular marker can reliably distinguish true RGCs from transient intermediates or off-target neurons with overlapping antigen expression. This limitation is particularly relevant for THY1-based enrichment strategies, which are commonly used for immunopanning and positive selection [9–12]. In our study, THY1 expression was variable and lacked specificity, being detected across multiple retinal and non-retinal neuronal populations. These findings suggest that THY1-based approaches are likely to isolate a heterogeneous mixture rather than a well-defined RGC population and that more stringent multi-marker or transcriptomically informed selection strategies will be necessary to obtain high-fidelity RGC enrichment. However, identifying surface markers that are both specific and stable across RGC subtypes remains an important and unresolved challenge.

Another feature of our dataset is the presence of *HOX*-enriched cell populations, particularly in hESC-derived organoids. From a developmental perspective, this is unexpected, as the retina originates from the anterior neuroectoderm, whereas *HOX* genes pattern posterior regions of the CNS [76,77]. However, our findings are consistent with previous studies reporting upregulation of *HOX* gene programs in hPSC-derived retinal organoids [18,78,79]. Although the abundance and composition of these clusters vary across studies, the recurrent detection of posterior *HOX*-expressing cells suggests that off-target neural identities represent a common feature of current retinal organoid differentiation protocols. Modulating retinoid acid availability during early differentiation such as the use of B27 supplement without vitamin A can shift neural identity toward anterior domains [80,81], and could potentially reduce the proportion of *HOX*-enriched cells.

While this protocol yielded variable proportions of RGCs across samples, the resulting RGC content (19-45%), excluding the poorly differentiated retinal organoids from WAB_RGC1, is comparable to previously reported yields from hPSC-to-RGC differentiation strategies validated by scRNA-seq. Earlier studies reported RGC proportions of around 12% using a 2D differentiation protocol and 17% in RGC-enriched retinal organoid systems [18,83], indicating that the overall efficiency observed here is above the range of transcriptomically validated approaches. Nevertheless, sample-to-sample variability indicates opportunities for further optimisation, as early exclusion of poorly patterned organoids and refinement of dissociation timing may reduce retention of retinal progenitor and photoreceptor-committed cells [14,84].

## Limitations

A limitation of this study is that RGC subtype classification is based on transcriptional profiles rather than functional properties. Classical RGC subtypes are defined by morphology, connectivity and electrophysiological behaviour [33], which cannot be resolved by scRNA-seq alone. As a result, transcriptionally defined clusters may not map directly onto established functional RGC classes and may instead reflect developmental states, stress responses or culture-induced divergence. Future studies integrating transcriptomics with electrophysiology, connectivity mapping and functional assays will be required to directly molecular identity with RGC function.

## Data availability

All data have been deposited in the ArrayExpress database (https://www.ebi.ac.uk/arrayexpress).

## Supplementary Information

Nine supplementary figures, three supplementary tables, and an additional Excel file containing differential gene expression results with an adjusted *p*-value < 0.05.

## Acknowledgments

The authors acknowledge Dr. Magdaline Sakkas of the Melbourne Cytometry Platform, The University of Melbourne, for her instruction on using the Beckman Coulter CytoFLEX LX flow cytometer, and Dr. Peter Lau of the Australian Genome Research Facility for his assistance in processing samples for scRNA-seq analysis.

## Authors’ Contributions

Conceptualisation: all authors; Methodology: all authors; Investigation: J.Y.W.M.; Data analysis: J.Y.W.M.; Writing original draft: J.Y.W.M., A.P.; Writing review & editing, all authors; Funding Acquisition: all authors; Supervision and project administration: A.P.

## Funding

This research was supported by funding from PYC therapeutics, a University of Melbourne Department of Anatomy and Physiology Early Career Seeding Grants (JYWM) and a Dame Kate Campbell Fellowship (AP).

## Materials & Correspondence

Requests for materials and correspondence should be addressed to Jessica Ma and Alice Pébay.

## Competing Interests

A.P. is a scientific advisor to PYC Therapeutics, and a director and shareholder of CellTellus Pty Ltd.

