## Supplementary for "Single-cell analysis reveals cellular heterogeneity and limits of marker-based assessment in retinal ganglion cell-enriched organoid cultures"

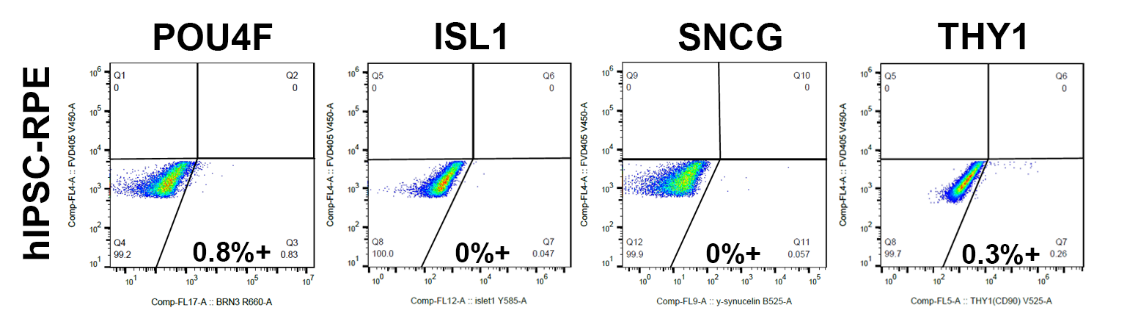

**Figure S1. Negative control for flow cytometry assessment.** hPSC-derived RPE was used as a negative control for the antibodies POU4F, ISL1, SNCG, and THY1.

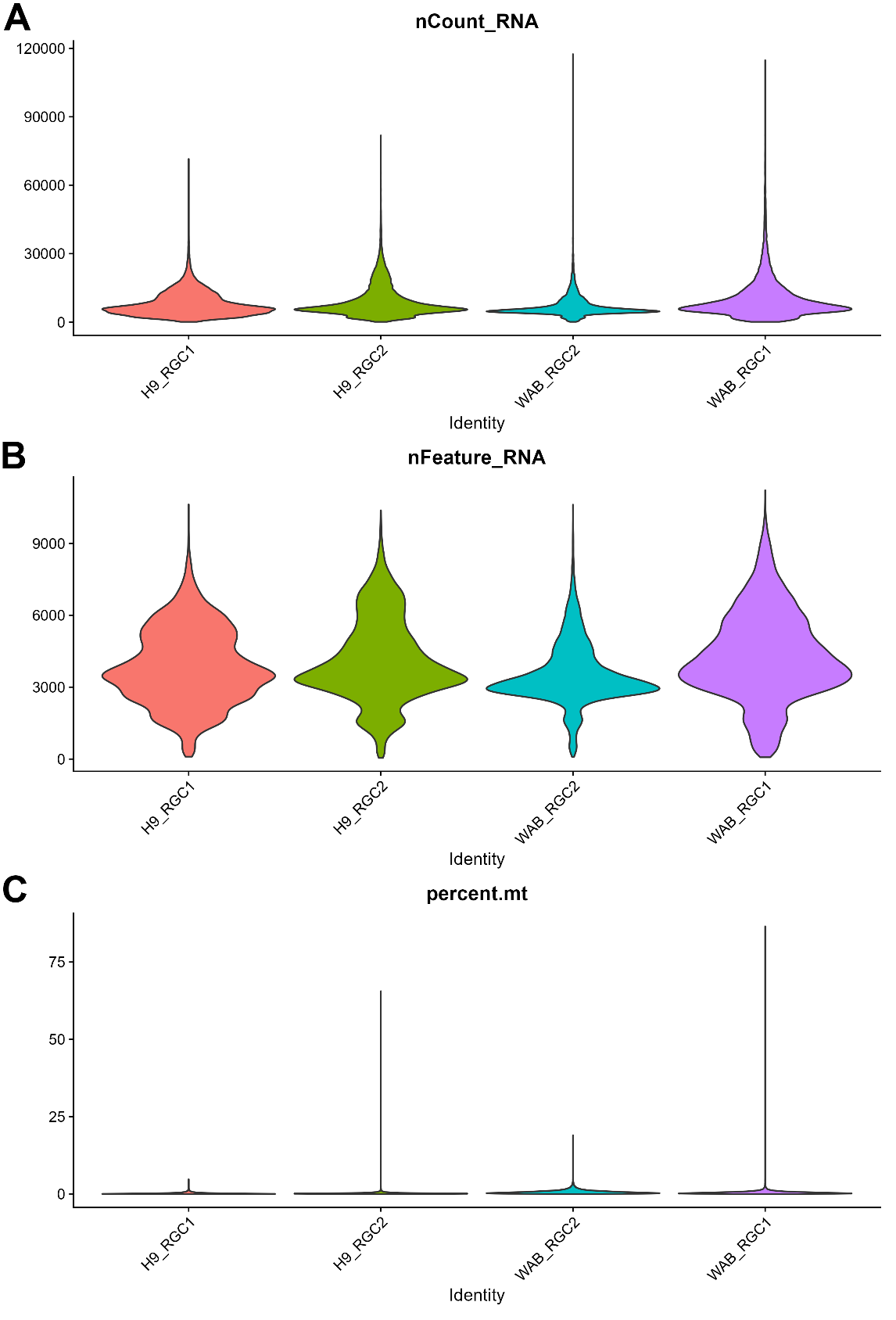

**Figure S2. Pre-filtering quality-control metrics.** Violin plots showing **(A)** total RNA counts per cell (nCount_RNA), showing the distribution of UMI counts across samples. **(B)** Number of detected genes per cell (nFeature_RNA). **(C)** Percentage of mitochondrial transcripts (percent.mt).

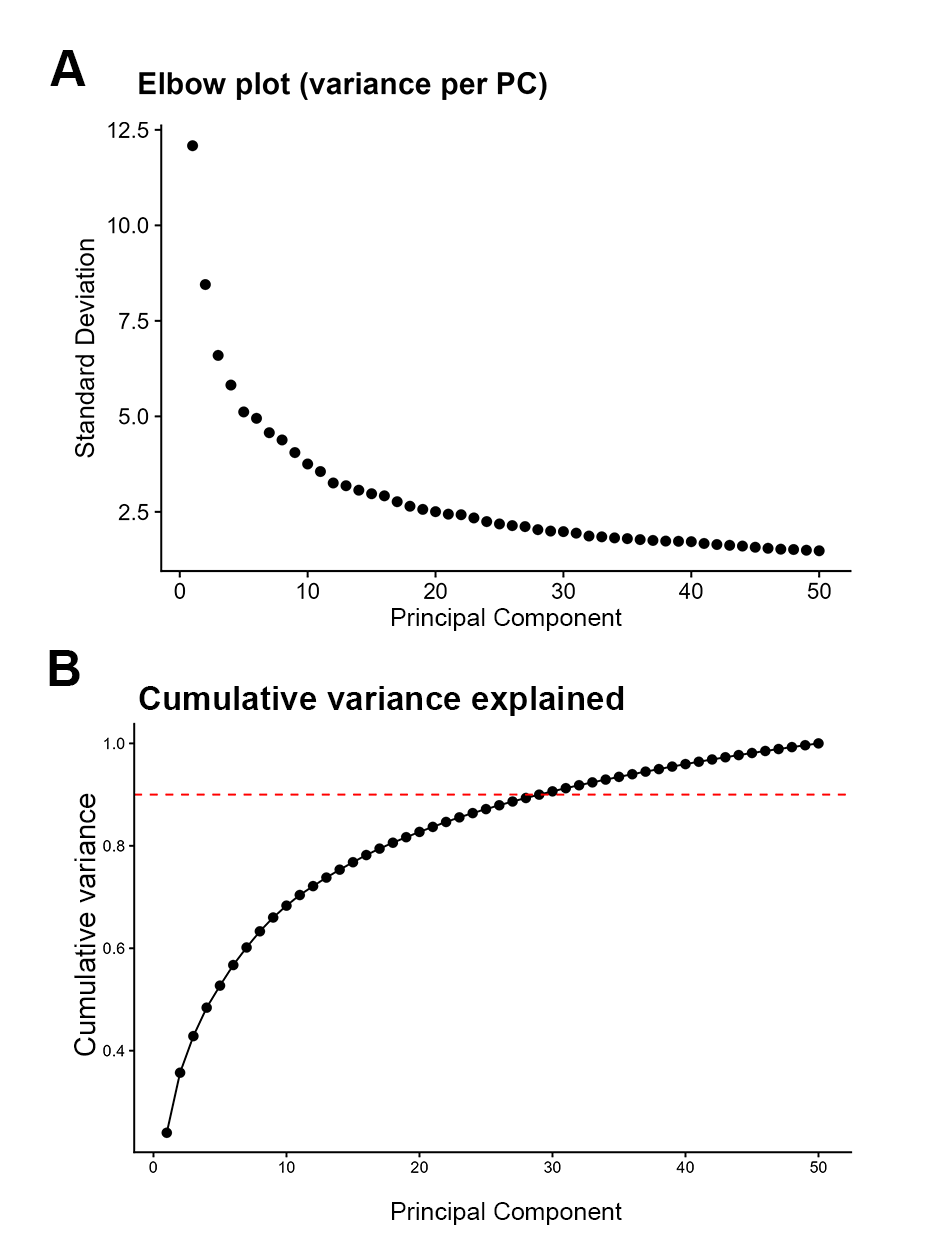

**Figure S3. Principal component selection for downstream single-cell analysis** (**A**) Elbow plot showing the point at which additional principal components (PCs) contribute minimal explanatory power. An inflection is evident at around PCs 20-25, indicating that the dominant biological structure is captured within this range. (**B**) Cumulative variance explained across PCs, with the 90% cumulative variance threshold reached at approximately PC30 (dashed line).

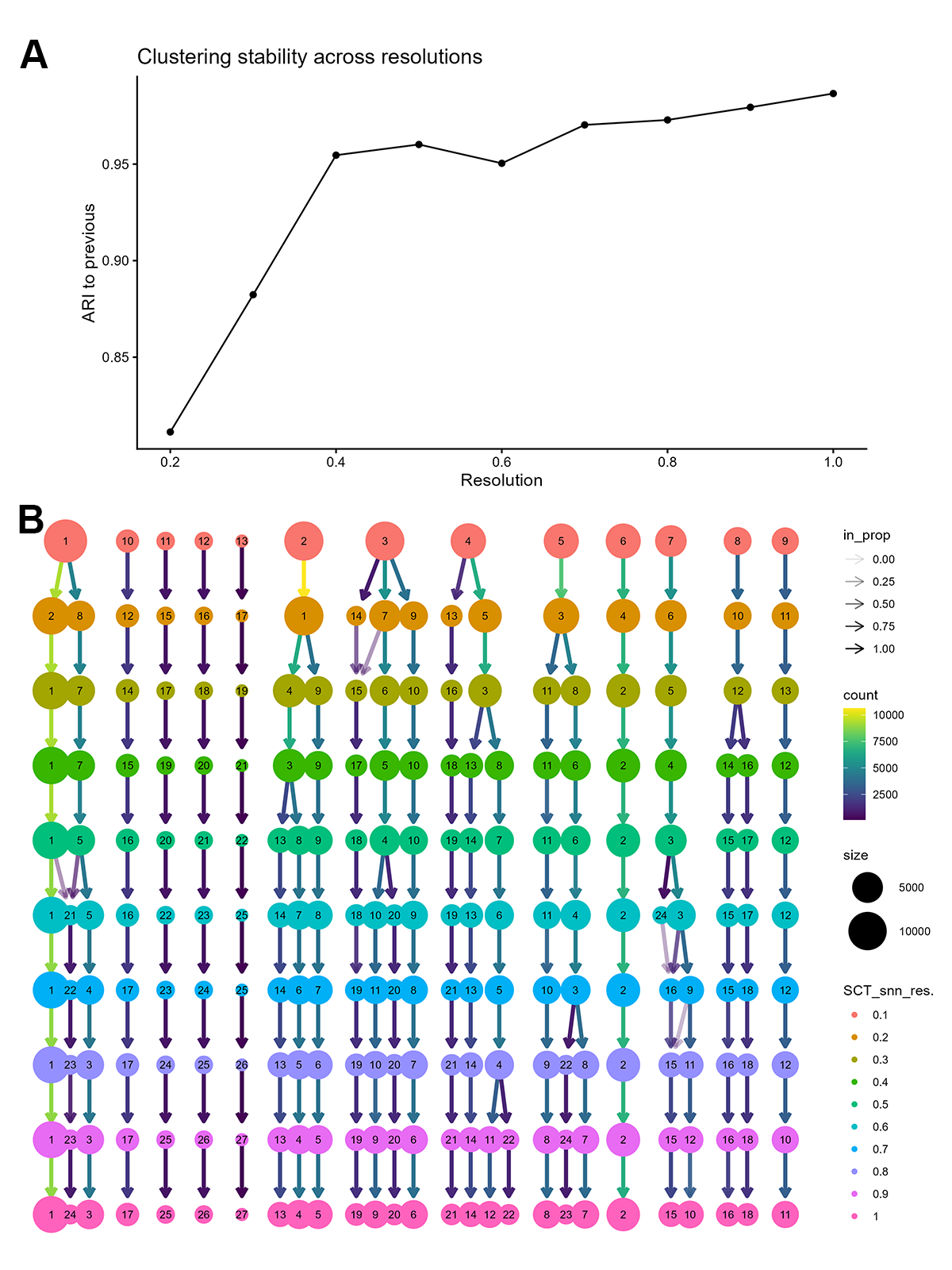

**Figure S4. Determination of clustering resolution based on ARI profiles and clustree analysis.** (**A**) Adjusted Rand Index (ARI) relative to the previous resolution, showing that cluster assignments stabilise at intermediate resolutions, with a clear plateau around resolution 0.5. (**B**) Clustree visualisation illustrating cluster propagation and stability across increasing resolutions across resolutions.

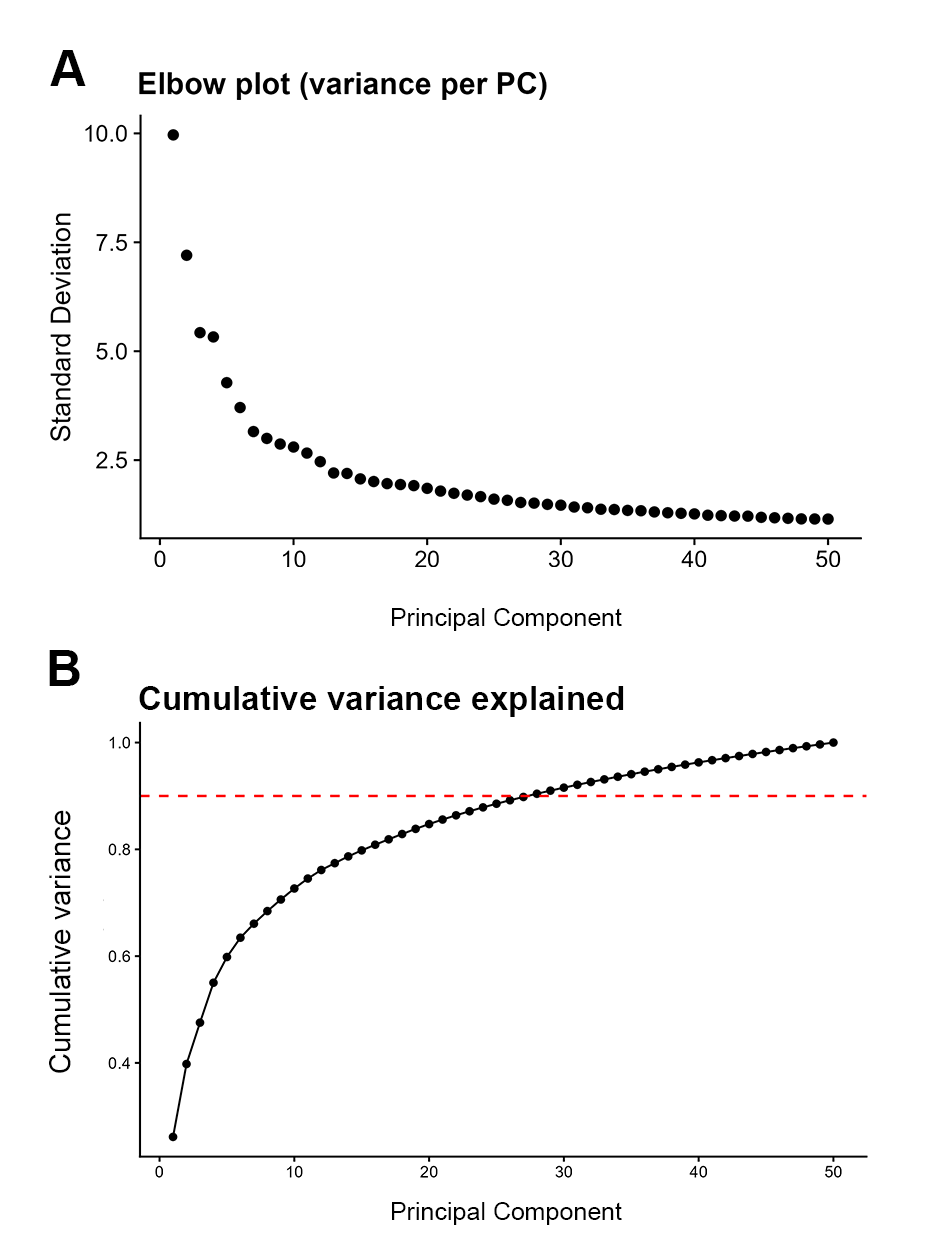

**Figure S5. Principal component selection for subclustering of RGC populations.** (**A**) Elbow plot showing the point at which additional principal components (PCs) contribute diminishing explanatory power, with an inflection observed around PC15. (**B**) Cumulative variance explained across PCs. Dashed horizontal lines indicate the 90% cumulative variance threshold used to guide PC selection.

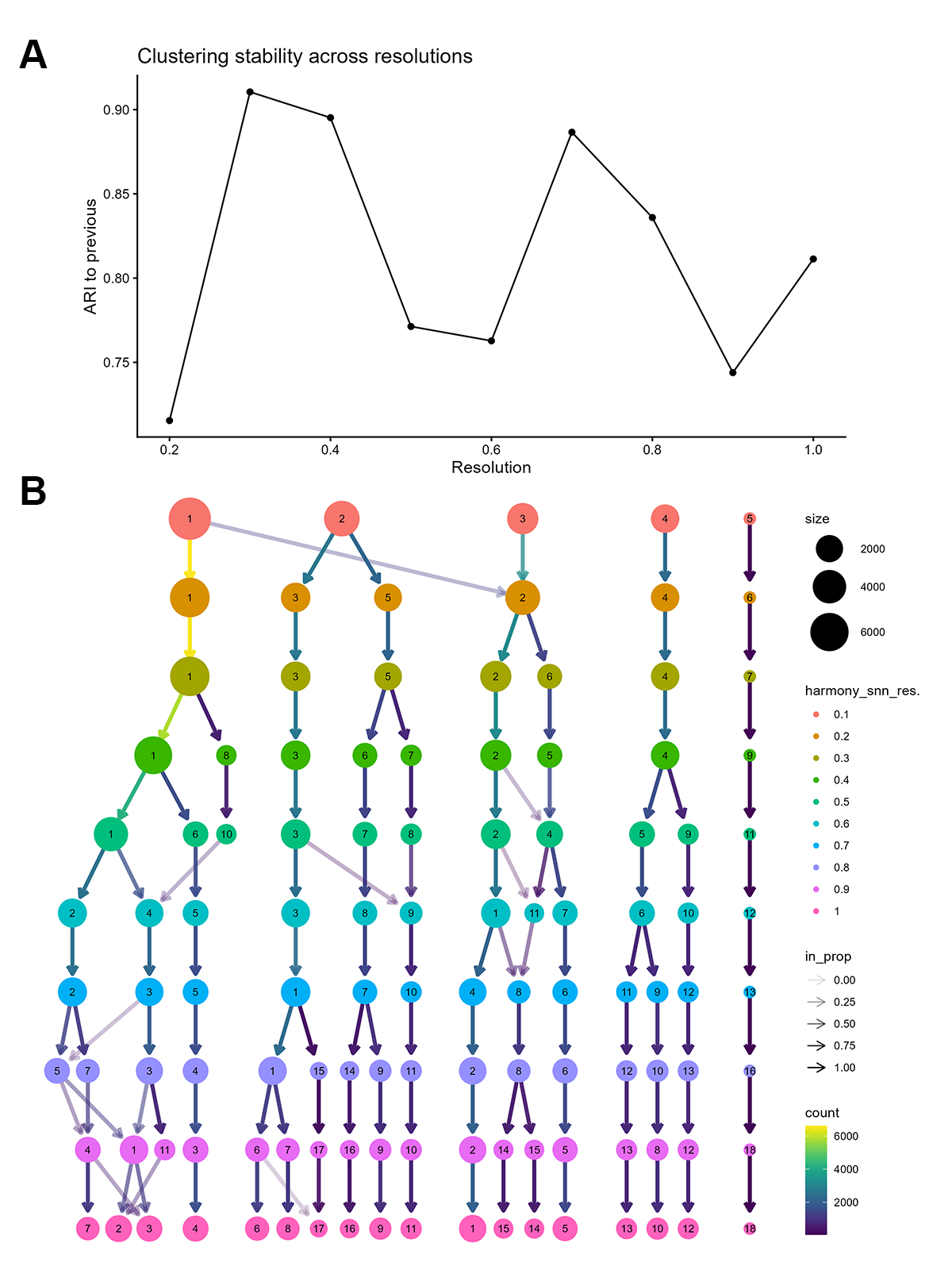

**Figure S6. Determination of clustering resolution based on Adjusted Rand Index (ARI) profiles and clustree analysis for the subclustering of RGC populations. (A)** The ARI-to-previous plot shows that cluster assignments reach their highest stability at around resolution 0.3. (B) Clustree visualisation illustrating cluster propagation across resolutions.

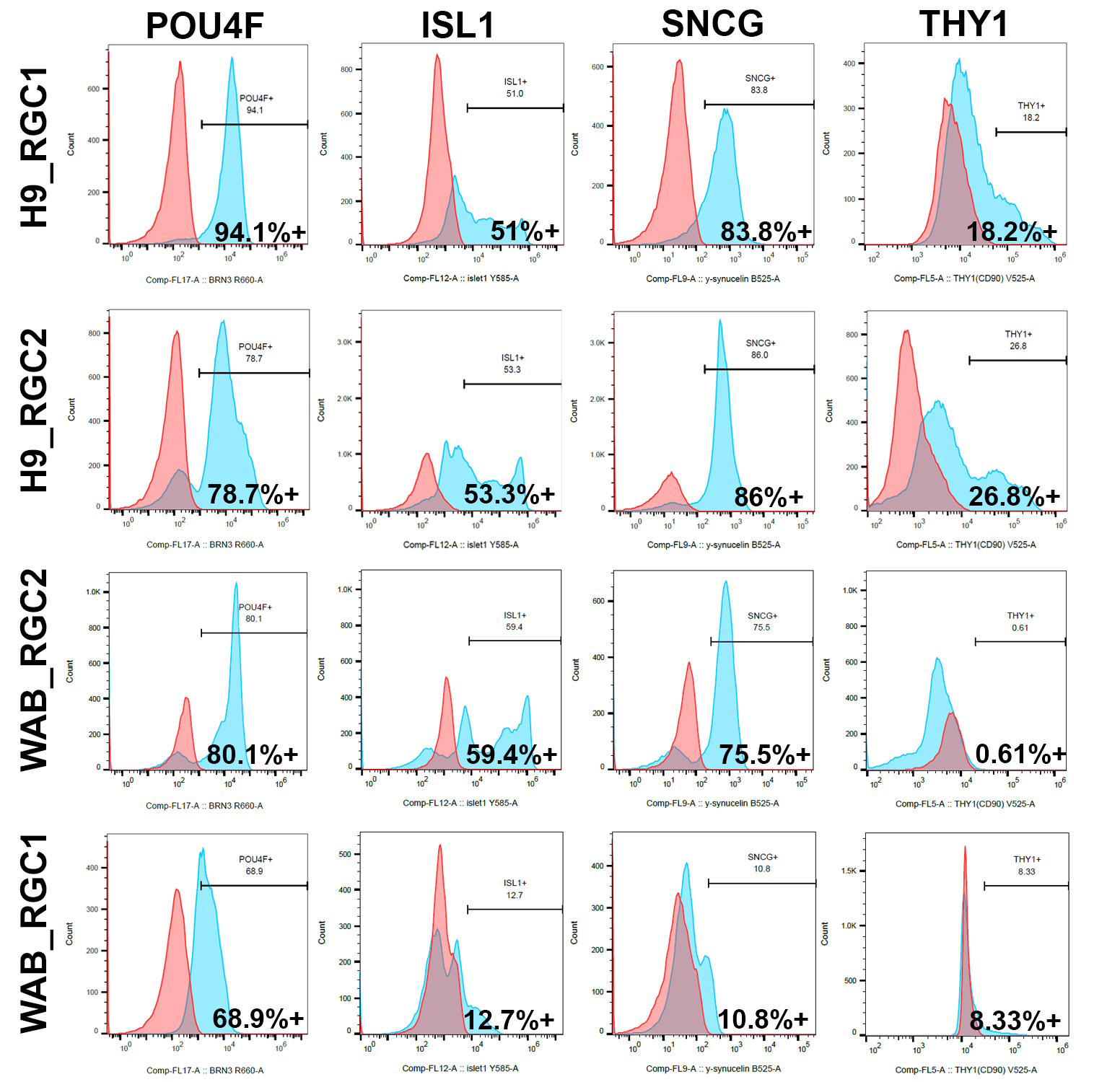

**Figure S7. Histogram representation of RGC marker expression measured by flow cytometry.** Representative fluorescence intensity histograms showing protein-level expression of POU4F, ISL1, SNCG, and THY1 across H9_RGC1, H9_RGC2, WAB_RGC2, and WAB_RGC1. Histograms depict the distribution of signal intensity within the live singlet population, with marker-positive fractions defined relative to unstained controls. Percentages indicate the proportion of marker-positive cells for each sample and marker.

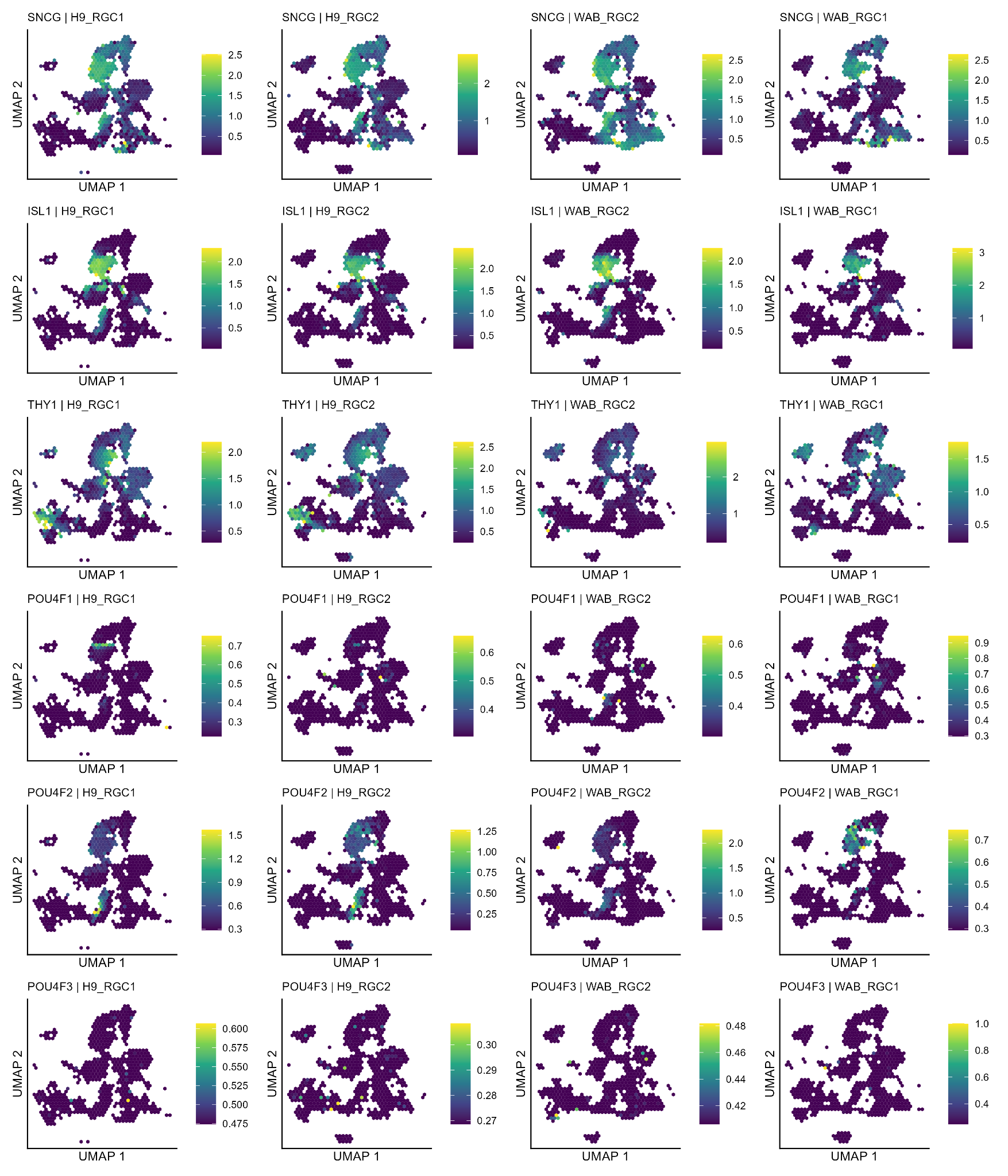

**Figure S8. Spatial distribution of RGC marker expression visualised using hexagon-binned UMAPs (schex).** Hexagon-binned UMAP visualisations showing mean log-normalised RNA expression of canonical RGC markers across the integrated scRNA-seq dataset, displayed separately for each sample (H9_RGC1, H9_RGC2, WAB_RGC2, and WAB_RGC1). Cells were aggregated into hexagonal bins on the UMAP embedding, and colour intensity represents the mean log-normalised expression across cells within each bin.

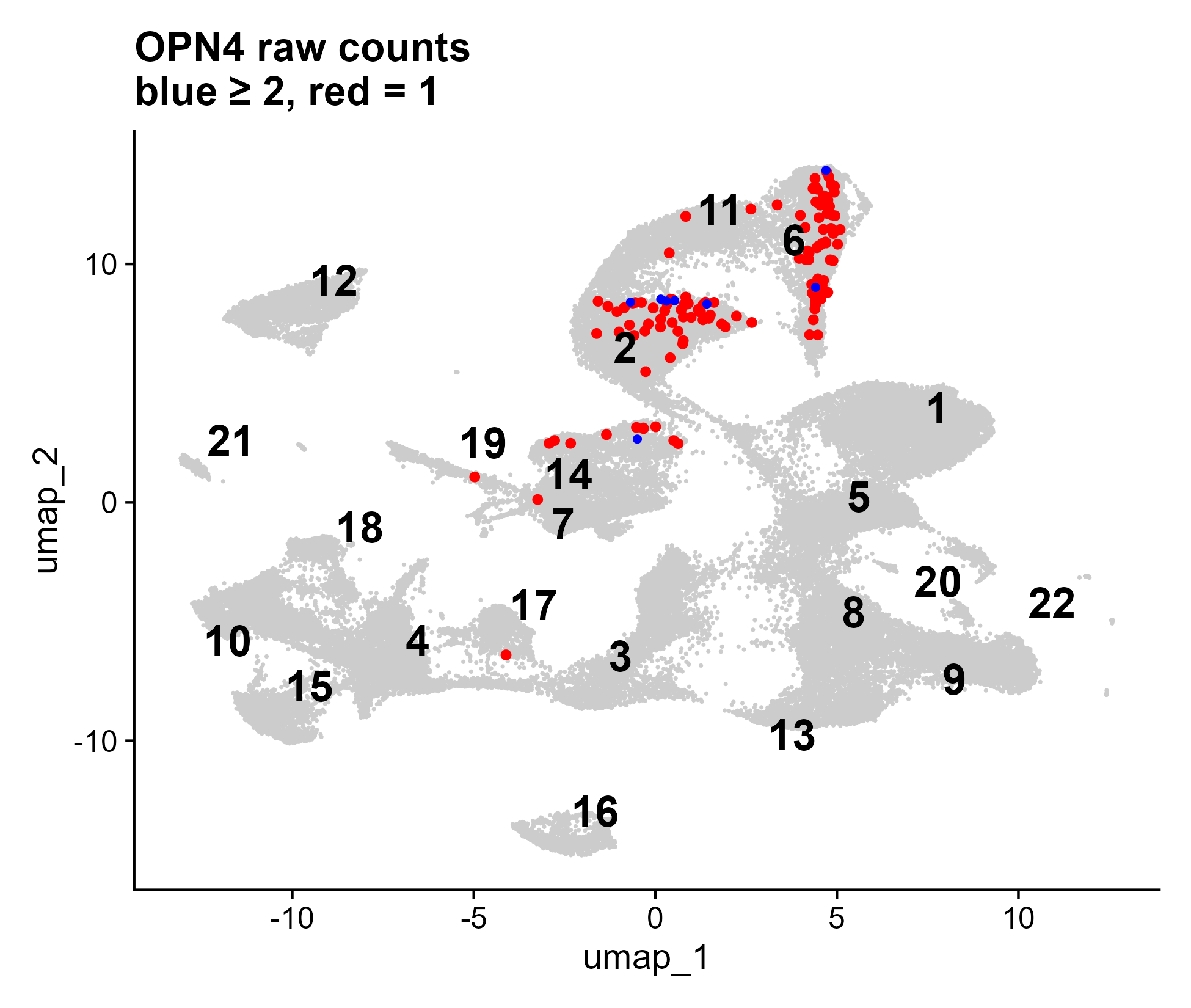

**Figure S9. UMAP visualisation of OPN4-expressing cells in the full integrated RGC-enriched retinal organoid dataset.** Each point represents a single cell; all cells are shown in grey, while OPN4-expressing cells are highlighted. Cells with low OPN4 abundance (SoupX-adjusted RNA count > 0 and < 2) are shown in red, and cells with higher abundance (SoupX-adjusted RNA count ≥ 2) are shown in blue, reflecting relative transcript abundance prior to normalisation.

**Table S1. Summary of cell retention across quality control steps*.*** Number of cells retained after EmptyDrops, SoupX + QC filtering, and doublet removal for each sample, along with the proportion of cells removed at each step.

| **Sample** | **Cells after EmptyDrops** | **Cells After SoupX+QC** | **Cells After Doublet** | **Integrated cells** |
| --- | --- | --- | --- | --- |
| **H9_RGC1** | **19,334** | **18,847** | **16,638** | **16,638** |
| **H9_RGC2** | **23,881** | **23,067** | **19,861** | **19,861** |
| **WAB_RGC2** | **21,678** | **21,140** | **18,926** | **18,926** |
| **WAB_RGC1** | **21,452** | **19,952** | **18,217** | **18,217** |

**Table S2. Distribution of identified cell types across individual datasets and clusters.** Per-cluster cell identities and distribution of cells across H9_RGC1, H9_RGC2, WAB_RGC2, and WAB_RGC1, with number and percentage of cells per dataset. Cluster annotations were assigned based on transcriptional profiles, including canonical markers.

|  |  | H9_RGC1 | | H9_RGC2 | | WAB_RGC2 | | WAB_RGC1 | |
| --- | --- | --- | --- | --- | --- | --- | --- | --- | --- |
| Cluster | Cell identity | # of cells | % of total | # of cells | % of total | # of cells | % of total | # of cells | % of total |
| 1 | Other (HOX-enriched) | 2,127 | 12.79 | 5,947 | 29.96 | 217 | 1.15 | 991 | 5.44 |
| 2 | RGC | 3,560 | 21.4 | 2,047 | 10.31 | 834 | 4.42 | 432 | 2.37 |
| 3 | RGC | 1,086 | 6.53 | 672 | 3.39 | 2,762 | 14.64 | 940 | 5.16 |
| 4 | Retinal progenitor cell | 1,278 | 7.68 | 920 | 4.63 | 487 | 2.58 | 2,247 | 12.35 |
| 5 | Amacrine cell | 1,036 | 6.23 | 1,797 | 9.05 | 682 | 3.61 | 1,339 | 7.36 |
| 6 | Horizontal cell | 1,160 | 6.97 | 688 | 3.47 | 2,271 | 12.04 | 506 | 2.78 |
| 7 | Amacrine cell | 1,495 | 8.99 | 1,886 | 9.5 | 364 | 1.93 | 539 | 2.96 |
| 8 | Photoreceptor- committed cell | 388 | 2.33 | 687 | 3.46 | 2,552 | 13.53 | 531 | 2.92 |
| 9 | Photoreceptor- committed cell | 168 | 1.01 | 1,133 | 5.71 | 2,146 | 11.37 | 661 | 3.63 |
| 10 | Retinal progenitor cell | 462 | 2.78 | 651 | 3.28 | 111 | 0.59 | 2,798 | 15.37 |
| 11 | RGC | 1,346 | 8.09 | 787 | 3.96 | 1,245 | 6.6 | 303 | 1.66 |
| 12 | Other | 21 | 0.13 | 818 | 4.12 | 1,447 | 7.67 | 948 | 5.21 |
| 13 | Photoreceptor- committed cell | 84 | 0.51 | 116 | 0.58 | 2,101 | 11.14 | 94 | 0.52 |
| 14 | RGC | 1,576 | 9.48 | 289 | 1.46 | 345 | 1.83 | 153 | 0.84 |
| 15 | Retinal progenitor cell | 201 | 1.21 | 145 | 0.73 | 392 | 2.08 | 1,230 | 6.76 |
| 16 | RPE | 2 | 0.01 | 81 | 0.41 | 30 | 0.16 | 1,678 | 9.22 |
| 17 | Retinal progenitor cell | 117 | 0.7 | 118 | 0.59 | 263 | 1.39 | 1,104 | 6.07 |
| 18 | Retinal progenitor cell | 184 | 1.11 | 419 | 2.11 | 82 | 0.43 | 698 | 3.83 |
| 19 | Multilineage (stressed) | 276 | 1.66 | 567 | 2.86 | 55 | 0.29 | 472 | 2.59 |
| 20 | Photoreceptor-committed cell | 35 | 0.21 | 72 | 0.36 | 464 | 2.46 | 22 | 0.12 |
| 21 | Retinal progenitor cell | 31 | 0.19 | 9 | 0.05 | 18 | 0.1 | 515 | 2.83 |

**Table S3. Distribution of *OPN4*-expressing cells across clusters in the full integrated RGC-enriched retinal organoid dataset.** Distribution of cells with detectable *OPN4* expression (raw count > 0) across clusters in the unnormalised, SoupX-adjusted RNA count matrix (n = 73,642). The analysis was conducted prior to RGC-specific subclustering and includes all major retinal and off-target cell populations.

| Cluster ID | Total OPN4+ cells | Count = 1 | Count ≥ 2 |
| --- | --- | --- | --- |
| 2 | 51 | 46 | 5 |
| 6 | 60 | 58 | 2 |
| 7 | 1 | 1 | 0 |
| 11 | 3 | 3 | 0 |
| 14 | 11 | 10 | 1 |
| 17 | 1 | 1 | 0 |
| 19 | 1 | 1 | 0 |
| Total | 128 | 120 | 8 |
